# Associations between physical activity and cognitive dysfunction in older companion dogs: Results from the Dog Aging Project

**DOI:** 10.1101/2022.04.20.488879

**Authors:** Emily E. Bray, David A. Raichlen, Kiersten K. Forsyth, Daniel E.L. Promislow, Gene E. Alexander, Evan L. MacLean, Dog Aging Project Consortium

**Author notes:** Contributed as senior authors (Gene E. Alexander, Evan L. MacLean).

## Abstract

Canine Cognitive Dysfunction (CCD) is a form of dementia that shares many similarities with Alzheimer’s disease. Given that physical activity is believed to reduce risk of Alzheimer’s disease in humans, we explored the association between physical activity and cognitive health in a cohort of companion dogs, aged 6-18 years. We hypothesized that higher levels of physical activity would be associated with lower (i.e., better) scores on a cognitive dysfunction rating instrument and lower prevalence of dementia, and that this association would be robust when controlling for age, comorbidities, and other potential confounders. Our sample included 11,574 companion dogs enrolled through the Dog Aging Project, of whom 287 had scores over the clinical threshold for CCD. In this observational, cross-sectional study, we used owner-reported questionnaire data to quantify dog cognitive health (via a validated scale), physical activity levels, health conditions, training history, and dietary supplements. We fit regression models with measures of cognitive health as the outcome, and physical activity—with several important covariates—as predictors. We found a significant negative relationship between physical activity and current severity of cognitive dysfunction symptoms (estimate = -0.10, 95% CI: -0.11 to - 0.08, *p* < 0.001), extent of symptom worsening over a 6-month interval (estimate = -0.07, 95% CI: -0.09 to -0.05, *p* <0.001), and whether a dog reached a clinical level of CCD (odds ratio = 0.53, 95% CI: 0.45 to 0.63, *p* <0.001). Physical activity was robustly associated with better cognitive outcomes in dogs. Our findings illustrate the value of companion dogs as a model for investigating relationships between physical activity and cognitive aging, including aspects of dementia that may have translational potential for Alzheimer’s disease. While the current study represents an important first step in identifying a relationship between physical activity and cognitive function, it cannot determine causality. Future studies are needed to rule out reverse causation by following the same dogs prospectively over time, and to evaluate causality by administering physical-activity interventions.

## Introduction

Alzheimer’s disease is a devastating, age-related progressive neurodegenerative brain disorder that leads to cognitive decline and dementia. It is therefore a high priority for researchers to identify early, modifiable risk factors that can be targeted as interventions (Raichlen & Alexander, 2017; Yu et al., 2020). Over the past few decades, physical activity has emerged as one such factor that may play an important role in reducing the risk of Alzheimer’s disease. There is evidence in humans that engaging in physical activity can have protective effects on cognitive function (Ahlskog, Geda, Graff-Radford, & Petersen, 2011; Santos-Lozano et al., 2016). In one large interventional study of adults with memory impairment, participating in a physical activity program for six months led to measurable increases in cognitive performance over the next year and a half (Lautenschlager et al., 2008). In a different intervention, researchers documented an increase in hippocampal volume linked to aerobic exercise training (Erickson et al., 2011). A meta-analysis across 12 cohorts including thousands of participants also concluded that physical activity significantly protected against cognitive decline, even at low to moderate levels (Sofi et al., 2011). A recent study found that late-life physical activity was associated with higher presynaptic protein levels, known to positively affect cognition (Casaletto et al., 2021). Indeed, recent meta-analyses of randomized controlled trials using physical activity interventions reveal notable protective effects for dementia risk (Beckett, Ardern, & Rotondi, 2015; Xu et al., 2017).

Several nonhuman species have been used as animal models for the cognitive impairments associated with Alzheimer’s disease (Cotman & Berchtold, 2007). Similarly to the human studies, there is preliminary evidence from work in rodents (Berchtold, Castello, & Cotman, 2010; Jahangiri, Gholamnezhad, & Hosseini, 2019; Van Praag, Shubert, Zhao, & Gage, 2005) and primates (Rhyu et al., 2010) that exercise enhances cognitive function and leads to neurogenesis, potentially protecting against the development of dementia. However, current model systems have limited translational potential due to reliance on genetically homogenous populations studied in artificial environments. To date, most comparative studies have been conducted using transgenic mouse models that attempt to mimic specific aspects of Alzheimer’s disease neuropathology, including the pathological deposition of amyloid-β (Aβ) plaques and neurofibrillary tangles with hyperphosphorylated tau (Jankowsky & Zheng, 2017). However, these models have typically focused on the least prevalent form in humans (Webster, Bachstetter, Nelson, Schmitt, & Van Eldik, 2014). No mouse model exhibits the full progression of Alzheimer’s disease, and the supraphysiological overexpression of amyloid precursor protein transgenes may alter brain development in ways that limit translational potential (Elder, Gama Sosa, & De Gasperi, 2010). In addition, studies with laboratory mice have limited ability to model the complex gene × environment interactions believed to underlie the heterogeneity observed in the development and progression of Alzheimer’s disease (Chouliaras et al., 2010).

Companion dogs have been proposed as a model for aging research with high translational potential (Creevy, Akey, Kaeberlein, & Promislow, 2022; Kaeberlein, Creevy, & Promislow, 2016). Unlike laboratory populations, companion dogs are genetically heterogeneous, and share many important features with humans, including the same living environments, disease risks and burdens, patterns of actuarial aging, and access to a sophisticated health care system (Hoffman, Creevy, Franks, O’Neill, & Promislow, 2018). Dogs have also been suggested as a valuable natural complementary model for the age-related dementia of Alzheimer’s disease. With advanced age, many dogs spontaneously develop a range of cognitive and behavioral impairments that resemble those associated with brain aging and Alzheimer’s dementia. Dozens of studies have shown that signs of age-related neurodegeneration in dogs are often accompanied by cognitive dysfunction in learning and memory analogous to impairments often seen in aging and Alzheimer’s disease (Head, 2011, 2013; Milgram et al., 2004; Packer et al., 2018; Ruehl et al., 1995). Although the full complement of Alzheimer’s disease neuropathology has yet to be consistently observed in any naturally occurring non-human animal model, Alzheimer-like pathology, e.g., Aβ 1-42, increases with age in companion dogs (Urfer et al., 2021) and has been described in the context of diffuse plaque deposition that has been related to cognitive decrements in older dogs (Cotman and Head, 2008). There is also preliminary evidence for tauopathy, another feature of Alzheimer-like pathology, in the brains of dogs diagnosed with canine cognitive dysfunction (Abey et al., 2021).

In addition, similarly to humans, physical activity as part of enrichment programs in dogs has been associated with reductions in Aβ Alzheimer-like pathology and improved cognitive performance (Cotman & Berchtold, 2007). Despite the strong potential for dog models of Alzheimer’s disease, most studies to date have used small laboratory samples that do not capitalize on the many potential benefits of a companion dog model (e.g., large heterogeneous populations living in the same environments as humans).

Previous exploratory work has looked broadly for associations between a wide range of characteristics and Canine Cognitive Dysfunction, finding that age as well as a single rating of physical activity were associated with Canine Cognitive Dysfunction (Yarborough, 2021). Building upon these findings, in the current observational study we focused our investigation on the relationship between physical activity and age-related impairments in cognitive function in companion dogs, using questionnaire data generated by The Dog Aging Project. Specifically, owners were asked to report the dog’s lifestyle (not active to active) as well as the typical duration and intensity of their dog’s physical activity. This dataset was analyzed alongside the owners’ responses to a validated instrument (Salvin, McGreevy, Sachdev, & Valenzuela, 2011) assessing behaviors indicative of cognitive dysfunction and dementia (i.e., changes in social activity; challenges in navigation, searching, and recognition). We hypothesized that higher levels of physical activity would be associated with lower (i.e., better) scores on a cognitive dysfunction rating instrument, and decreased risk of dementia, and that this association would be robust when controlling for age, comorbidities, and potential confounders (e.g., joint supplements, motor impairments, exercise intolerance). Additionally, given that we know little about potential risk factors and protective effects for canine dementia, we also examined associations between several lifestyle factors (i.e., use of neuroprotective supplements and engagement in formal dog training activities) and categories of health conditions (i.e., neurologic conditions, sensory deficits, periodontal disease, and liver failure) with dementia outcomes.

## Methods

### Subjects

All dogs were members of the Dog Aging Project (DAP), a nationwide research study of companion dogs that aims to better understand the biological and environmental factors that impact health span and lifespan (Creevy et al., 2022; Kaeberlein et al., 2016). While the DAP is an ongoing longitudinal study, the data in the current study were cross-sectional, drawing on initial responses from owners whose dogs are enrolled in the first cohort. Owners completed the requested online surveys between December 26, 2019 and December 31, 2020 (Dog Aging Project, 2021). Study data were collected and managed using REDCap electronic data capture tools hosted through the DAP (Harris et al., 2019; Harris et al., 2009). These data are publicly available and housed on the Terra platform at the Broad Institute of MIT and Harvard.

### Instruments

Upon enrollment in the DAP, owners completed the Health and Life Experience Survey (HLES). In addition to collecting dog and owner demographics, this detailed questionnaire also asked owners to report on their dog’s physical activity, environment, behavior, diet, medications and preventatives, and health status. For the current study, we were mainly interested in the data reflecting physical activity and health status.

After completing HLES, all participants were asked to participate in a second survey: the Canine Social and Learned Behavior Survey (CSLB). The intent of this survey was to measure owner-report of cognitive function. The CSLB, renamed by the DAP, is based on the Canine Cognitive Dysfunction Rating Scale (CCDR) (Salvin et al., 2011), with minor wording modifications to select items. The CCDR was presented to participants as the Canine Social and Learned Behavior Survey to avoid the negative connotations of the phrase ‘cognitive dysfunction’. This instrument asks owners to indicate the frequency with which their dogs exhibit behaviors indicative of cognitive dysfunction and dementia (i.e., disengagement from social activity; difficulty in navigation, searching, and recognition). Based on owner responses, dogs receive a score that ranges from 16 to 80, where higher scores are indicative of worse cognitive function. This instrument was previously validated in a sample of dogs 8 years and older as a way of distinguishing dogs with CCD from those without (Salvin et al., 2011). In the current manuscript, we also explored its utility as a continuous measure.

During the study period, we received HLES responses from 27,541 unique DAP participants, of which 20,096 went on to also complete a CSLB.

### Ethical Note

The University of Washington IRB deemed that recruitment of dog owners for the DAP, and the administration and content of the DAP HLES, are human subjects research that qualifies for Category 2 exempt status (IRB ID no. 5988, effective 10/30/2018). No interactions between researchers and privately owned dogs occurred; therefore, IACUC oversight was not required.

### Inclusion/Exclusion Criteria

Given that cognitive decline is not typically observed in dogs until at least six years of age (Harvey, 2021; Packer et al., 2018; Studzinski et al., 2006), we specified age of inclusion as 6 ≤ age < 18 years at the time of CSLB completion.

After applying this exclusion criterion, the final sample consisted of 11,574 dogs whose owners completed both the HLES and CSLB surveys. CSLB was always completed at least one week after completion of HLES. Most participants in the final sample (87.8%) completed CSLB within 3 months of completing HLES and always within one year (range: 7 to 352 days, mean: 47.14 days).

### Outcome variable

Our outcome of interest was the owner-reported symptoms of cognitive dysfunction of each dog, which we measured via three scores derived from CSLB responses. We first performed principal component analysis (PCA) on the 13 response items (see SI 1, Appendix A for survey questions). Parallel analysis recommended retaining two principal components. We used an oblimin rotation to allow correlation between the two PCs (see Table S1 in SI 1 for loadings). The first PC, which we called ‘change’, was loaded highly by questions regarding reported changes in cognitive dysfunction symptoms over the prior 6 months. The second PC, which we called ‘severity’, was loaded highly by items measuring reported current symptom severity. Finally, we analyzed Canine Cognitive Dysfunction (CCD) status as a binary exposure, wherein dogs who scored 50 or above were deemed to be above the diagnostic clinical threshold for CCD, and dogs below this score were not (Salvin et al., 2011).

### Predictor Variables

Our main predictor of interest was physical activity. To calculate this variable for each dog, we performed PCA on three HLES-reported activity variables: lifestyle activity level (reported as not active, moderately active, or very active over the past year), average activity intensity level (reported as low: walking, medium: jogging, or vigorous: sprinting, such as playing fetch or frisbee), and average daily time spent physically active (reported in hours and minutes). Parallel analysis recommended retaining one principal component from these measures. This principal component explained 52% of the variance and was loaded positively by all three questions regarding physical activity. We used the scores from this component as our measure of physical activity (PA-score). Initial exploratory analyses suggested substantial and linear declines in physical activity with age (Fig S1 in SI 1).

We used information reported in HLES about diverse medical conditions with potential to influence cognitive function or physical activity level as covariates. Specifically, based on past literature, we expected the following health-related factors to be associated with risk of cognitive impairment in dogs: neurologic conditions, such as epilepsy (Hobbs et al., 2020; Watson, Packer, Rusbridge, & Volk, 2020; Winter, Packer, & Volk, 2018), sensory deficits in the visual and auditory domains (Fischer et al., 2016; Ford et al., 2018; Szabó, Miklósi, & Kubinyi, 2018), periodontal disease (Dewey & Rishniw, 2021; Harding, Gonder, Robinson, Crean, & Singhrao, 2017; Singhrao, Harding, Poole, Kesavalu, & Crean, 2015), and liver failure (Butterworth, 2016; Felipo, 2013).

We also created covariates for orthopedic conditions and exercise intolerance, which we expected to be negatively associated with physical activity levels. In the exercise intolerance category, we accounted for cardiac and respiratory conditions that negatively affect a dog’s ability to exercise—either by rendering the dogs unable to exert themselves physically, or because the prevailing veterinary advice for the diagnosis is restricted activity.

Lastly, to control for other factors potentially influencing general health, we created variables for whether dogs had been diagnosed with certain systemic disorders, including cancer and those affecting the kidneys and the endocrine system.

For each of the health condition categories described above, all participants were assigned a binary score (affected/unaffected). Dogs were considered ‘affected’ if their owner reported them to have one or more relevant conditions within a given category. We only included chronic conditions that were likely to affect the relevant systems, and thus excluded temporary conditions that, given standard recommended medical care, would only temporarily affect the relevant systems. For example, in the orthopedic category, we scored hip dysplasia as an ‘affected’ condition, as it is a long-term issue that affects mobility, whereas fractured bones were not included because the most likely prognosis is complete recovery and therefore the impact on physical activity is temporary. For cataracts and ligament ruptures, we only included dogs as affected (in the sensory impairment and orthopedic categories, respectively) if the diagnosis was *not* followed by surgery. Our curated list of health conditions included in each covariate category can be found in SI 2, and the full list of health conditions that owners were asked about is listed in SI 3.

Additionally, we created covariates for lifestyle factors that preliminary evidence suggests might have ameliorating or protective effects for physical activity and/or cognition. If dogs received glucosamine and/or other joint supplements daily, they were considered ‘affected’ in the joint supplement category (McCarthy et al., 2007). If dogs received omega 3, vitamins, probiotics, antioxidants, taurine, carnitine, and/or coenzyme Q10 daily, they were considered ‘affected’ in the neuroprotective supplement category (Heath, Barabas, & Craze, 2007; Mad’ari, Farbakova, & Žilka, 2017; Milgram et al., 2004; Pan, Kennedy, Jönsson, & Milgram, 2018). Finally, we also created a variable accounting for whether a dog had a history of training (Bray et al., 2022), given intriguing preliminary evidence that this sort of enrichment is linked to delay in cognitive decline (Bray et al., 2022; Milgram, Siwak-Tapp, Araujo, & Head, 2006; Szabó et al., 2018). Training history was determined according to what the owner reported as the dog’s primary or secondary activity (e.g., service dogs, agility dogs, and dogs trained for field trials vs. pets/companion; see SI 1, Appendix B for full details).

A summary of the demographic variables, incidence of health conditions, physical activity levels, training history, and dietary supplement use within our sample is reported in Table 1, broken down by participants who met the diagnostic score for CCD (*n* = 287) and those who did not (*n* = 11,287).

**Table 1.**
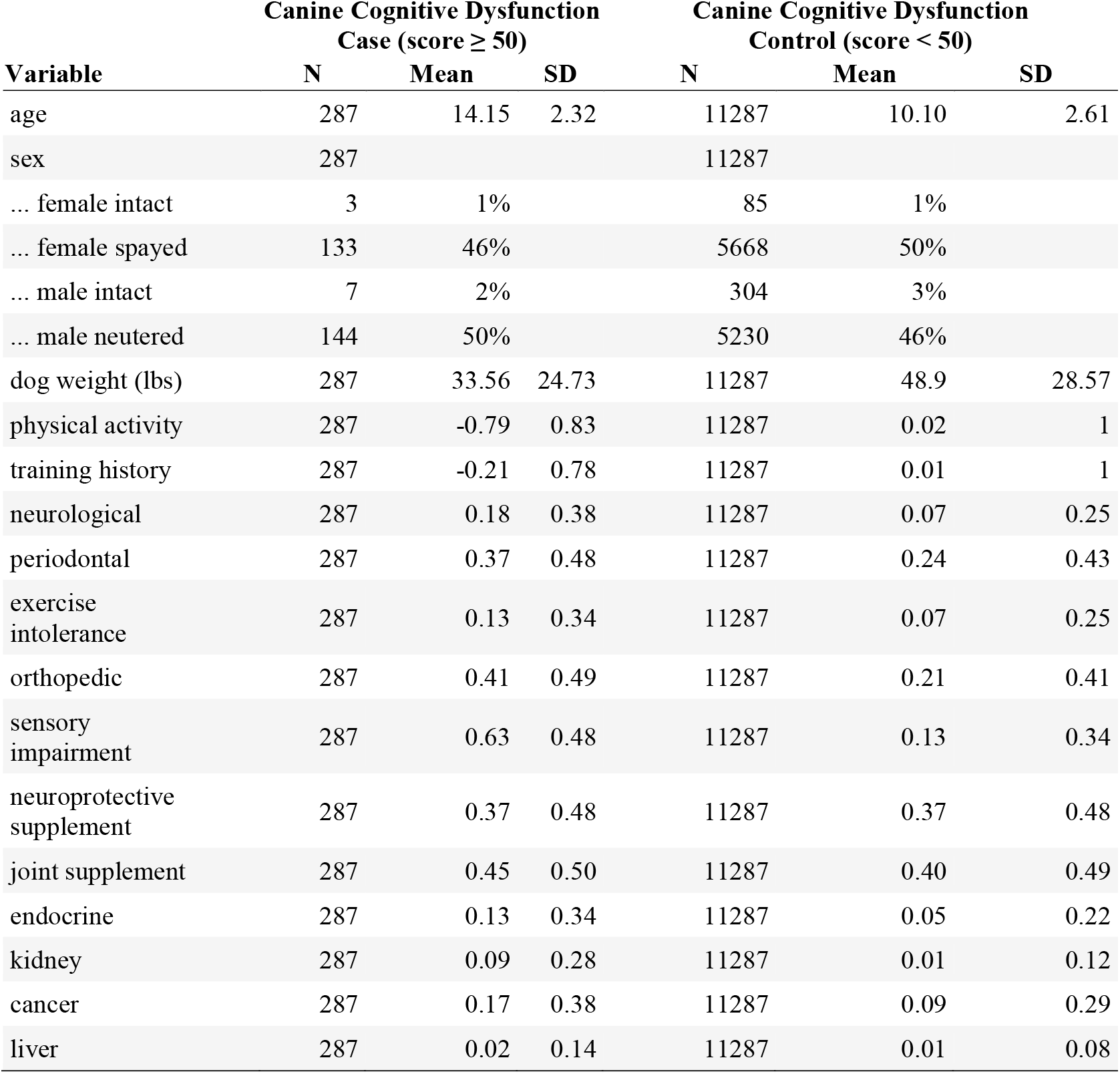
Summary statistics of our sample.

### Statistical Methods

All statistical analyses were carried out in R v.4.0.3 (R Development Core Team, 2016).

We fit three tiers of models for each of our outcome variables. In our first tier of analysis, we built a base model that included only key predictor variables (physical activity and age) and a minimal set of covariates. The effect of age was modelled using a second-order polynomial term because preliminary exploratory analyses revealed a non-linear relationship between age and the cognitive outcomes (see Fig S2 in SI 1). The other covariates included in our base models included dog sex (female, intact; female, spayed; male, intact; male, castrated), dog size (lbs), and owner age (18-24, 25-34, 35-44, 45-54, 55-64, 65-74, 75+). For models using the categorical measure of dementia status as the outcome, the owner age variable was collapsed to two levels (18-54, 55+) and dog sex was collapsed to two levels (male, female) to avoid small cell sizes.

In our second tier of analysis, we built a model that included all the variables from our base model as well as hypothesis-driven confounders and risk or protective factors. The additional variables for these models included whether a given dog exhibited sensory impairments (e.g., visual and/or auditory), motor impairments (e.g., orthopedic challenges), exercise intolerance (e.g., cardiac and/or respiratory challenges), neurological conditions other than dementia (i.e., dogs with a reported diagnosis of dementia or senility—and no other neurological conditions— were considered ‘unaffected’ in this category), periodontal disease, liver disease, as well as whether they were currently receiving joint and/or neuroprotective supplements, and whether they had a history of training. For models using the categorical measure of dementia as the outcome, liver disease was removed as a covariate due to small cell sizes when stratifying on this covariate.

Finally, in the third tier of analysis, we added the remaining, non-hypothesis driven covariates, for health condition categories including endocrine disease, kidney disease, and cancer.

We applied our three-tier modeling approach to the three different outcome variables, using linear regressions for symptom severity and recent symptom change, and a logistic regression for CCD status. Continuous outcomes (severity and change) were subjected to an inverse rank normal transformation to better meet the assumptions of linear modeling, and then standardized to have a mean of 0 and standard deviation of 1, to facilitate interpretation. We fit a total of nine statistical models (three for each dependent measure). To identify the best model for each outcome, we compared the Akaike information criterion scores across models.

We also performed some sensitivity analyses. To determine if any observed associations would still hold in a cognitively healthy population, we re-ran our original analyses but removed all dogs above the CCD threshold (*n* = 11,287, Tables S2 and S3 in SI 1). Given that over half of our sample was comprised of mixed breed dogs (*n* = 6,027 (52%)), a highly heterogenous group, we did not control for breed in our main analyses. Thus, in a follow-up set of sensitivity analyses, we first repeated all models but eliminated all purebred dogs from the sample (*n* = 6,027, Table S4-S6 in SI 1). Additionally, we then repeated all models but only included purebred dogs—using breeds with at least 10 dogs in the dataset (*n* = 5,167 dogs from 92 breeds; Table S7 in SI 1), and, for the CCD model, at least one member of the breed above the CCD threshold (*n* = 3,945 dogs from 53 breeds; Table S8 in SI 1)—and added breed as a covariate (Table S4-S6 in SI 1). Finally, based on the possibility that CSLB scores below 20 may be implausible, we re-ran the models from our main analyses, excluding the subset of dogs with a score of 19 and lower (*n* = 11,368; see SI 1 for details).

## Results

For all outcomes, results from each of the three tiers of analysis displayed the same pattern but the fully adjusted model fit the best in all cases, as assessed by the lowest Akaike information criterion (Tables 2-4). Therefore, the results reported below are derived from the models including all candidate covariates.

**Table 2.** Model comparisons between the three tiers of models predicting symptom severity, reporting the beta coefficients and the 95% confidence interval based on robust standard errors in parentheses. Age effects are shown in Fig 1.

| Parameter | Symptom Severity |  |  |  |  |  |
| --- | --- | --- | --- | --- | --- | --- |
|  | Minimally Adjusted |  | Moderately Adjusted |  | Fully Adjusted |  |
|  | Beta (95% CI)<sup>1</sup> | p-value | Beta (95% CI)<sup>1</sup> | p-value | Beta (95% CI)<sup>1</sup> | p-value |
| physical activity | -0.116 (-0.134 to -0.098) | <0.001 | -0.096 (-0.114 to -0.079) | <0.001 | -0.095 (-0.113 to -0.077) | <0.001 |
| dog weight (lbs) | 0.000 (-0.001 to 0.000) | 0.123 | 0.000 (-0.001 to 0.001) | 0.720 | 0.000 (-0.001 to 0.001) | 0.941 |
| sex |  |  |  |  |  |  |
| female intact | — |  | — |  | — |  |
| female spayed | 0.238 (0.056 to 0.419) | 0.010 | 0.198 (0.020 to 0.376) | 0.029 | 0.195 (0.018 to 0.373) | 0.031 |
| male intact | 0.208 (0.004 to 0.411) | 0.046 | 0.171 (-0.029 to 0.372) | 0.093 | 0.170 (-0.030 to 0.370) | 0.096 |
| male neutered | 0.290 (0.108 to 0.471) | 0.002 | 0.244 (0.066 to 0.422) | 0.007 | 0.243 (0.066 to 0.420) | 0.007 |
| owner age |  |  |  |  |  |  |
| 18-24 | — |  | — |  | — |  |
| 25-34 | -0.348 (-0.567 to -0.129) | 0.002 | -0.336 (-0.545 to -0.127) | 0.002 | -0.334 (-0.542 to -0.125) | 0.002 |
| 35-44 | -0.578 (-0.794 to -0.362) | <0.001 | -0.556 (-0.762 to -0.350) | <0.001 | -0.554 (-0.759 to -0.348) | <0.001 |
| 45-54 | -0.725 (-0.939 to -0.510) | <0.001 | -0.710 (-0.915 to -0.505) | <0.001 | -0.707 (-0.911 to -0.503) | <0.001 |
| 55-64 | -0.876 (-1.09 to -0.664) | <0.001 | -0.861 (-1.06 to -0.658) | <0.001 | -0.856 (-1.06 to -0.654) | <0.001 |
| 65-74 | -0.99 (-1.20 to -0.774) | <0.001 | -0.97 (-1.18 to -0.772) | <0.001 | -0.97 (-1.17 to -0.767) | <0.001 |
| 75 and older | -1.07 (-1.29 to -0.852) | <0.001 | -1.05 (-1.26 to -0.844) | <0.001 | -1.05 (-1.25 to -0.839) | <0.001 |
| sensory impairment |  |  | 0.408 (0.351 to 0.464) | <0.001 | 0.405 (0.349 to 0.461) | <0.001 |
| orthopedic |  |  | 0.087 (0.044 to 0.130) | <0.001 | 0.084 (0.041 to 0.127) | <0.001 |
| exercise intolerance |  |  | 0.045 (-0.020 to 0.111) | 0.177 | 0.043 (-0.023 to 0.108) | 0.203 |
| neurological |  |  | 0.076 (0.008 to 0.143) | 0.028 | 0.073 (0.005 to 0.140) | 0.035 |
| periodontal |  |  | 0.063 (0.024 to 0.101) | 0.002 | 0.060 (0.021 to 0.099) | 0.003 |
| liver |  |  | 0.041 (-0.182 to 0.264) | 0.720 | 0.030 (-0.193 to 0.253) | 0.790 |
| joint supplement |  |  | -0.032 (-0.074 to 0.009) | 0.125 | -0.031 (-0.073 to 0.010) | 0.139 |
| neuroprotective supplement |  |  | -0.078 (-0.119 to -0.038) | <0.001 | -0.082 (-0.123 to -0.042) | <0.001 |
| training history |  |  | -0.031 (-0.047 to -0.014) | <0.001 | -0.031 (-0.047 to -0.014) | <0.001 |
| endocrine |  |  |  |  | 0.085 (0.009 to 0.161) | 0.029 |
| kidney |  |  |  |  | 0.112 (-0.025 to 0.248) | 0.109 |
| cancer |  |  |  |  | 0.057 (0.001 to 0.113) | 0.047 |
| AIC | 30,326 |  | 30,010 |  | 30,003 |  |
| <sup>l</sup> CI = Confidence Interval |  |  |  |  |  |  |

**Table 3.**
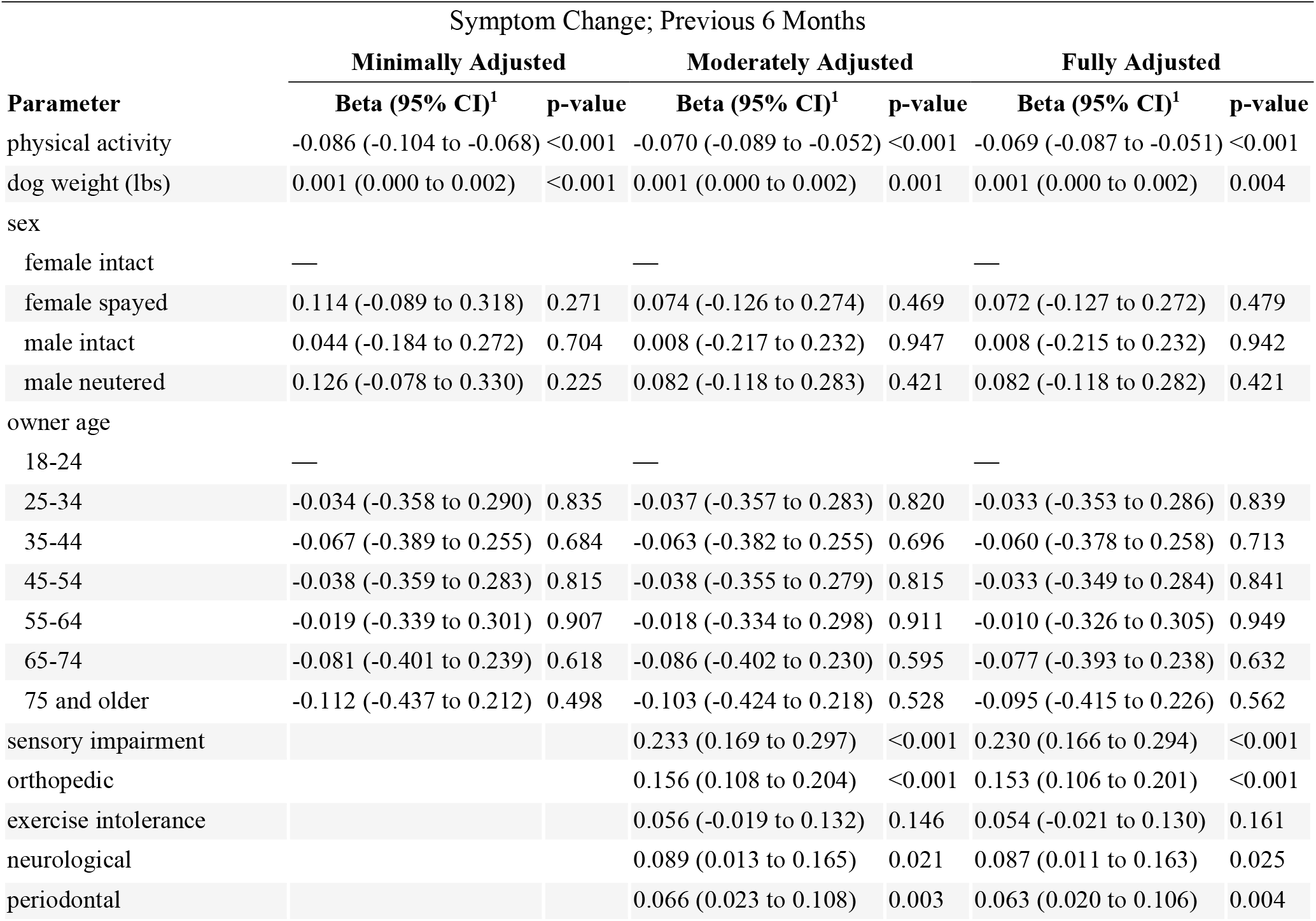
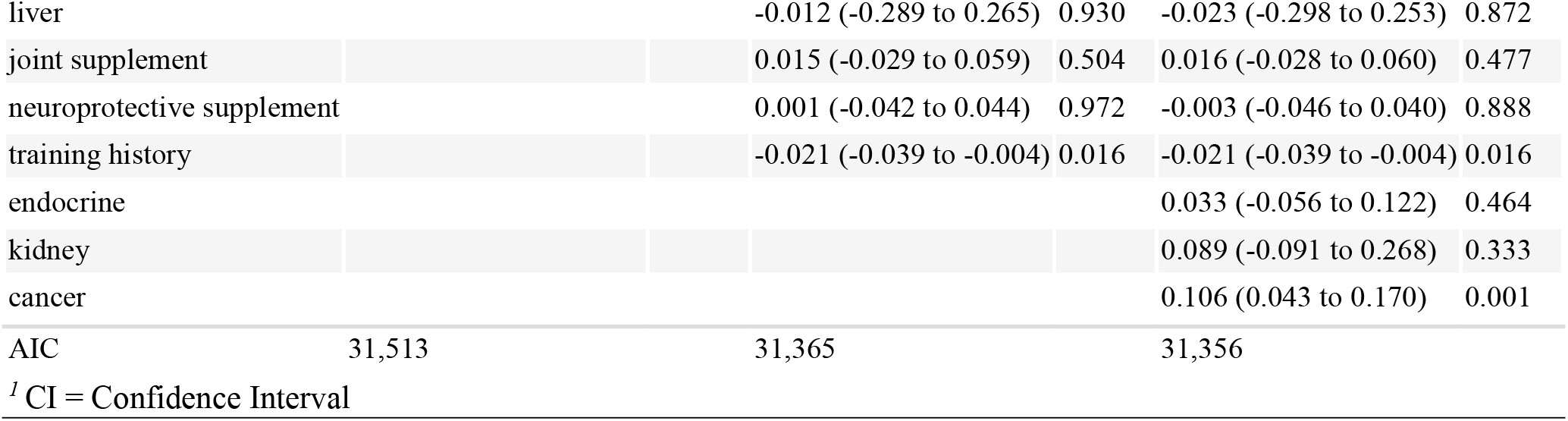
Model comparisons between the three tiers of models predicting cognitive decline in previous six months, reporting the beta coefficients and the 95% confidence interval based on robust standard errors in parentheses. Age effects are shown in Fig 1.

**Table 4.** Model comparisons between the three tiers of models predicting CCD status, reporting the odds ratio and the 95% confidence interval in parentheses. Age effects are shown in Fig 1.

|  | Canine Cognitive Dysfunction (Clinical Cutoff) |  |  |  |  |  |
| --- | --- | --- | --- | --- | --- | --- |
|  | Minimally Adjusted |  | Moderately Adjusted |  | Fully Adjusted |  |
| Parameter | OR (95% CI)<sup>1</sup> | p-value | OR (95% CI)<sup>1</sup> | p-value | OR (95% CI)<sup>1</sup> | p-value |
| physical activity | 0.51 (0.43 to 0.60) | <0.001 | 0.53 (0.45 to 0.62) | <0.001 | 0.53 (0.45 to 0.63) | <0.001 |
| dog weight (lbs) | 0.99 (0.99 to 1.00) | 0.003 | 0.99 (0.99 to 1.00) | 0.008 | 0.99 (0.99 to 1.00) | 0.006 |
| dog sex |  |  |  |  |  |  |
| male | — |  | — |  | — |  |
| female | 0.85 (0.66 to 1.09) | 0.198 | 0.85 (0.65 to 1.09) | 0.202 | 0.83 (0.64 to 1.07) | 0.152 |
| owner age |  |  |  |  |  |  |
| 18-54 | — |  | — |  | — |  |
| 55 and older | 0.78 (0.61 to 1.01) | 0.062 | 0.75 (0.58 to 0.97) | 0.029 | 0.78 (0.60 to 1.02) | 0.070 |
| sensory impairment |  |  | 3.23 (2.45 to 4.28) | <0.001 | 3.20 (2.43 to 4.24) | <0.001 |
| orthopedic |  |  | 1.22 (0.92 to 1.61) | 0.160 | 1.22 (0.92 to 1.62) | 0.162 |
| exercise intolerance |  |  | 0.98 (0.66 to 1.43) | 0.928 | 0.97 (0.65 to 1.42) | 0.887 |
| neurological |  |  | 1.31 (0.91 to 1.86) | 0.137 | 1.29 (0.89 to 1.84) | 0.162 |
| periodontal |  |  | 0.80 (0.60 to 1.05) | 0.105 | 0.78 (0.59 to 1.02) | 0.076 |
| joint supplement |  |  | 0.96 (0.70 to 1.32) | 0.822 | 1.00 (0.72 to 1.37) | 0.979 |
| neuroprotective supplement |  |  | 1.02 (0.74 to 1.40) | 0.898 | 0.97 (0.71 to 1.34) | 0.872 |
| training history |  |  | 0.89 (0.75 to 1.04) | 0.174 | 0.88 (0.74 to 1.03) | 0.133 |
| endocrine |  |  |  |  | 1.46 (0.97 to 2.16) | 0.062 |
| kidney |  |  |  |  | 1.85 (1.09 to 3.04) | 0.017 |
| cancer |  |  |  |  | 1.15 (0.80 to 1.61) | 0.437 |
| AIC | 1,977 |  | 1,911 |  | 1,908 |  |
| <sup>1</sup> OR = Odds Ratio, CI = Confidence Interval |  |  |  |  |  |  |

As expected, all three cognitive outcomes were negatively impacted by age, with effect of age increasing at older ages (Fig 1). In all models, there was also a significant relationship between physical activity and cognitive outcomes (Fig 2).

**Fig 1.**
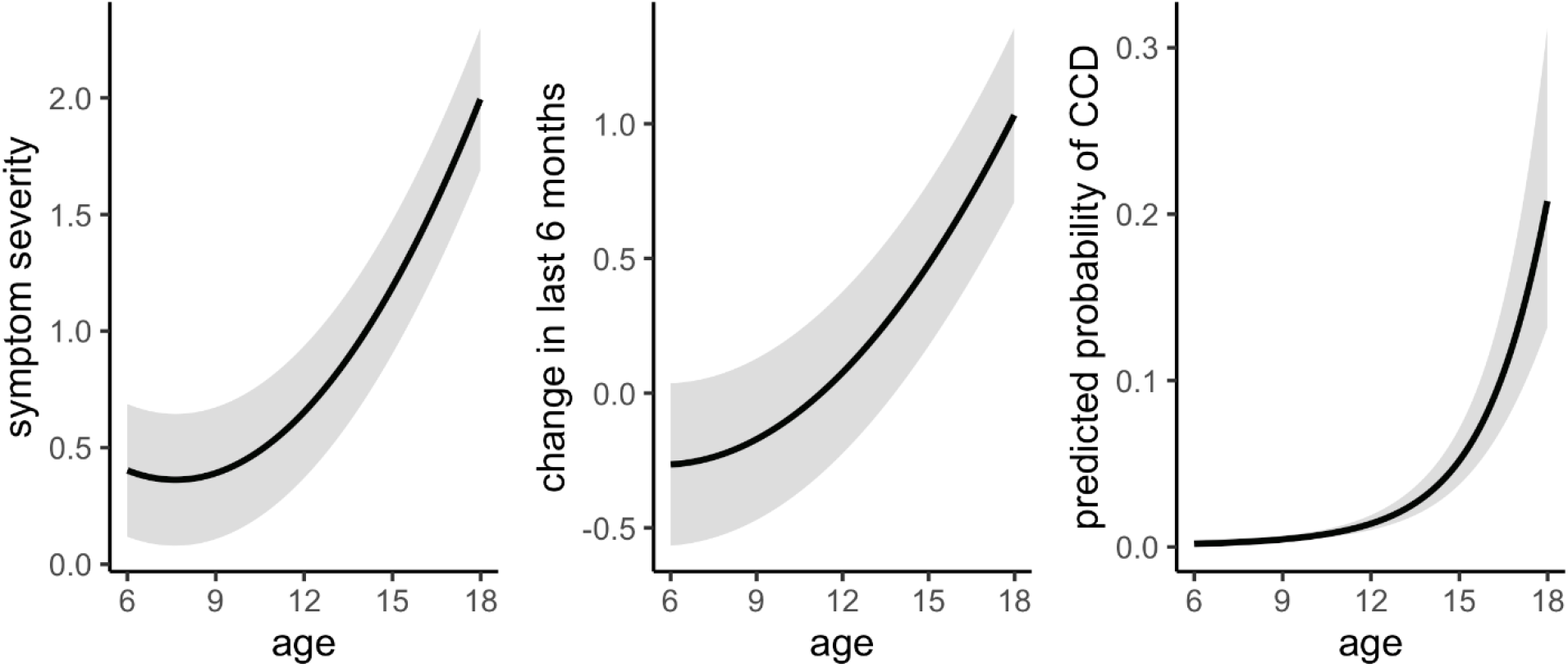
The estimated association between age and symptom severity (PCA-derived score), symptom change in last 6 months (PCA-derived score), and probability of a CCD diagnosis, respectively (with 95% confidence intervals indicated in gray). Results are from our fully adjusted models and include both linear and quadratic terms for age.

**Fig 2.**
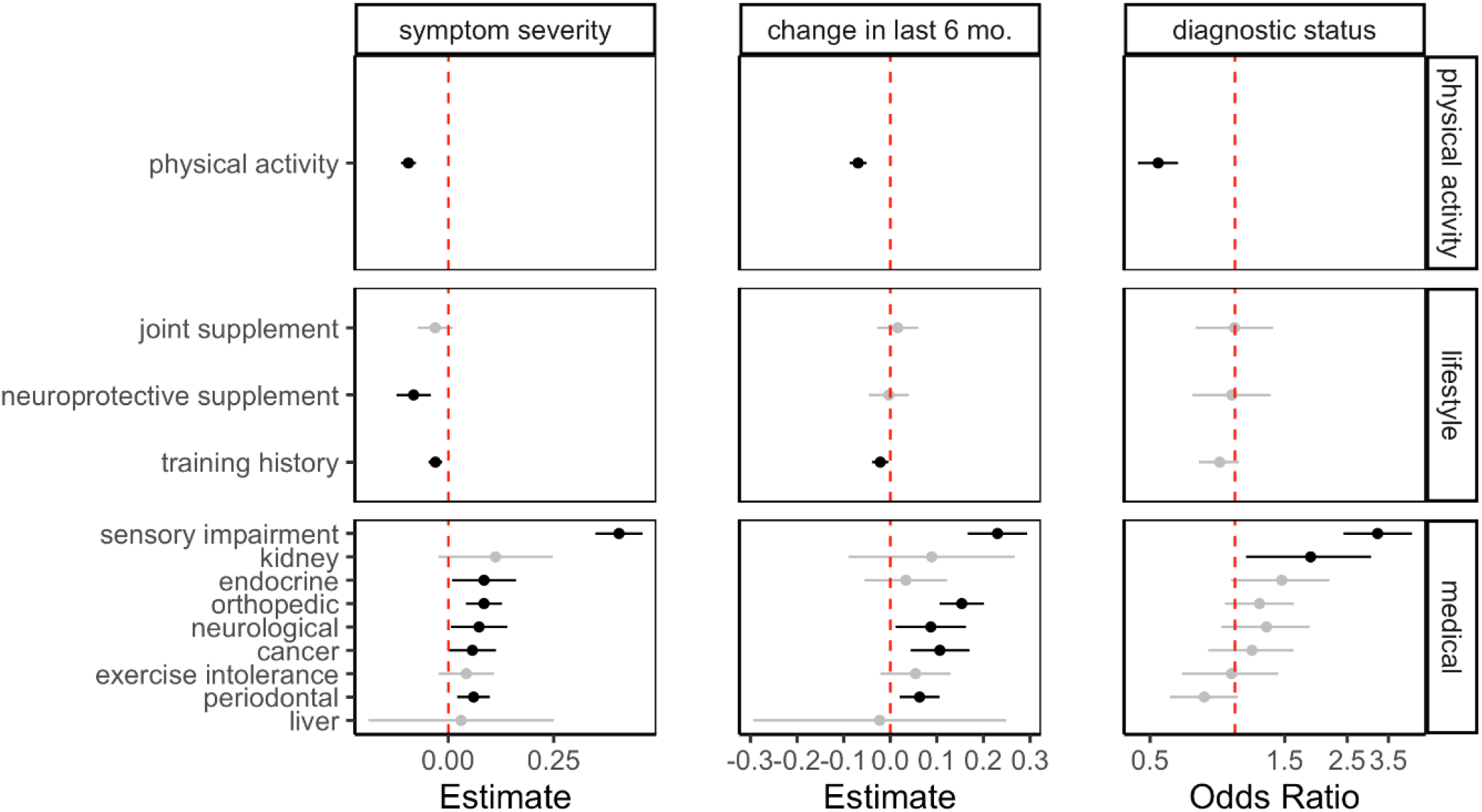
The beta coefficients (for the severity and change models) and odds ratios (for the CCD diagnosis model) of physical activity, as well as the other lifestyle (joint supplement, neuroprotective supplement, training history) and medical (sensory impairment, kidney, endocrine, orthopedic, neurological, cancer, liver, exercise intolerance, periodontal) covariates from the fully adjusted models. The red dotted line indicates the null expectation (i.e., 0 for the betas and 1 for the odds ratios). Significant findings are presented in black, while nonsignificant findings are presented in gray. The bars represent the 95% confidence intervals.

In the severity model, we found a significant negative association between physical activity and severity of cognitive symptoms, whereby high levels of activity were linked to lower (i.e., better) scores on the CSLB (Fig 2; Table 2). We also identified associations between two other hypothesized protective factors (training history and neuroprotective supplements), in which both a history of training and daily consumption of neuroprotective supplements were associated with better cognitive outcomes. For the final hypothesized protective factor (joint supplements), the beta coefficient was negative but not statistically significant. We also observed that poor health in certain domains was a risk factor for symptom severity. For our medical covariates, beta coefficients were positive and statistically significant for six categories of conditions (sensory impairment, endocrine, orthopedic, neurological, cancer, and periodontal) and positive but not statistically significant for the final three categories of conditions (kidney, liver, and exercise intolerance; Fig 2; Table 2). Results were similar in the analysis that excluded dogs above the CCD threshold (Table S2 in SI 1), suggesting that these relationships hold below the clinical cutoff for a diagnosis of dementia. Results were also similar in secondary analyses including only mixed breed dogs and dogs from the most common breeds (see Table S4 in SI 1). Across all three models, the negative association between symptom severity and our main exposure of interest (physical activity) remained significant, as did the negative associations with training history and neuroprotective supplements and the positive associations with two categories of medical conditions (sensory impairment and orthopedic). Finally, removing dogs with reported CSLB scores less than 20 did not change our findings (Table S9 in SI 1).

In the symptom change model, we again found a significant negative relationship between physical activity and reported change in cognitive symptoms as recalled by owners over the prior 6-month period, whereby higher levels of activity were linked to less owner-reported cognitive decline across the preceding six months (Fig 2; Table 3). We also identified a negative association with one of our other hypothesized protective factors (training history), in which dogs with an extensive training history exhibited less cognitive decline in the preceding six months. For the two other hypothesized protective factors (neuroprotective and joint supplements), the beta coefficients were near zero and not statistically significant. We also found evidence that poor health in certain domains was a risk factor for symptoms worsening over a 6-month period. For our medical covariates, beta coefficients were positive and statistically significant for five categories of medical conditions (sensory impairment, orthopedic, neurological, cancer, and periodontal), and not statistically significant for four categories of conditions (kidney, endocrine, exercise intolerance, and liver). Results were similar when performing our original analyses but removing all dogs above the CCD threshold (Table S3 in SI 1), suggesting that these relationships hold below the clinical cutoff for a diagnosis of dementia. Results were also similar in secondary analyses including only mixed breed dogs and dogs from the most common breeds (see Table S5 in SI 1): across all three models, the negative association between symptom change and physical activity remained significant, as did the positive associations with three categories of medical conditions (sensory impairment, orthopedic, and periodontal). Finally, removing dogs with reported CSLB scores less than 20 did not change our findings (Table S10 in SI 1).

In the CCD status model, we found that higher levels of physical activity were associated with lower odds of being over the diagnostic threshold for CCD (Fig 2; Table 4). The adjusted odds ratio was 0.53 (95% CI: 0.45 to 0.63) and statistically significant for physical activity, but there were no significant associations with the other hypothesized protective factors (training history, neuroprotective supplements, and joint supplements). We also found evidence that poor health in certain domains was associated with CCD, whereby individuals with CCD were also likely to have other owner-reported health issues. For our medical covariates, we observed OR > 1.0 and statistically significant for three categories of medical conditions (sensory impairment, kidney, and endocrine) with none of the other six categories of conditions (orthopedic, neurological, cancer, liver, exercise intolerance, and periodontal) reaching statistical significance. Results were similar in secondary analyses including only mixed breed dogs and dogs from the most common breeds (see Table S6 in SI 1 for full report): across all three models, the negative association between being over the diagnostic threshold for CCD and physical activity remained significant, as did the positive association with sensory impairment. Removing dogs with reported CSLB scores less than 20 did not change our findings (Table S11 in SI 1).

## Discussion

We investigated the relationship between physical activity and cognitive health in a sample of over 10,000 companion dogs. By exploring this relationship in a large population living in an environment shared with humans, we aimed to gain insight regarding factors associated with healthy cognitive aging and to identify potential modifiable risk factors that may prevent cognitive dysfunction and dementia (Deckers et al., 2015).

Across all models, we observed robust associations between physical activity and cognitive health. Physical activity was significantly negatively associated with three metrics of cognitive dysfunction: current symptom severity, extent of worsening over a 6-month interval, and whether a dog had reached a clinical threshold for CCD. These results held when controlling for basic demographic factors (weight, sex, and age of the dog, as well as age of the owner), hypothesis-driven confounders and risk factors related to lifestyle (joint-enhancing supplements, neuroprotective supplements, and training history) and health (sensory impairments, exercise intolerance, orthopedic conditions, neurological conditions other than dementia, periodontal disease, liver conditions), and other general health conditions (endocrine conditions, kidney failure, and cancer).

Furthermore, sensitivity analyses indicated that the association between physical activity and cognitive function held even when dogs who met the CCD threshold were removed from the sample. Thus, even in non-clinical cohorts physical activity may be associated with measurable cognitive benefits in older dogs, and/or declines in cognitive function may be associated with declines in owner-reported physical activity.

In addition to the association between physical activity and cognition, our analyses revealed relationships between cognitive health and several other health and lifestyle variables. For example, one of the strongest observed associations was between CSLB scores indicating worse cognitive health and sensory impairment, in line with the findings of a similar questionnaire-based study of 1,300 companion dogs (Szabó et al., 2018). While it may be that sensory impairment is a confounder (i.e., owners may mistakenly attribute a change in behavior to cognitive dysfunction when really it is the result of failing vision and/or audition), there is also evidence in the human literature that such impairments are potential risk factors for dementia (Hwang et al., 2020; Luo et al., 2018; Maharani, Dawes, Nazroo, Tampubolon, & Pendleton, 2020).

We also found a positive association between taking daily neuroprotective supplements (e.g., fish oil) and cognitive symptom severity. This finding is consistent with some clinical studies in dogs (Pan, Kennedy, et al., 2018; Pan, Landsberg, et al., 2018) and humans (Fotuhi, Mohassel, & Yaffe, 2009; Nolan, Mulcahy, Power, Moran, & Howard, 2018), although other studies in the human literature have found no effect (Danthiir et al., 2018; van de Rest et al., 2008). A potential limitation of this finding is that owners who are motivated to provide potentially neuroprotective supplements may be biased in their evaluation of their pet’s dementia symptoms. However, these supplements (e.g., fish oil) are also recommended by veterinarians for numerous other perceived benefits (e.g., heart health, coat shine, allergy relief, and pain management), so we do not know what expectations owners have regarding their potential effects on cognition.

Finally, we identified an association between two of our cognitive outcomes—symptom severity and cognitive change over the last 6 months—and training, whereby dogs who had a history of training were less likely to exhibit signs of cognitive decline. This finding is consistent with the idea that both physical exercise *and* mental exercise can have a beneficial impact on the brain (Marx, 2005; Raichlen & Alexander, 2017; Raichlen et al., 2020). Furthermore, this measure accounted for previous activity (i.e., history of training versus current training regimen) and so, given the timeline, cannot be readily explained by reverse causality. While the literature in humans (Kramer, Bherer, Colcombe, Dong, & Greenough, 2004) and laboratory animals (Birch & Kelly, 2019), including beagles (Milgram et al., 2005; Milgram et al., 2006), supports the idea that enrichment can lead to better cognitive functioning in old age, only one other study has demonstrated this relationship in companion dogs (Szabó et al., 2018). Nonetheless, this relationship has interesting potential parallels to associations between cognitive training and educational attainment in the context of dementia and Alzheimer’s disease risk in humans (Xu et al., 2016).

Our study has several notable limitations. First, despite the large sample size and wide range of covariates able to be accounted for, we cannot rule out unmeasured confounding. Second, all data were owner-reported and thus subject to potential pitfalls associated with self-report. Despite this limitation, the survey used in our analyses is known to have excellent diagnostic accuracy and test-retest reliability (Salvin et al., 2011). Third, we categorized dogs as either ‘affected’ or ‘not affected’ on each health covariate based on owner-reported diagnoses when filling out the HLES survey. However, HLES does not capture information about a condition’s severity. While all dogs were included in each category if they had a relevant diagnosis, in reality that condition might not have had a measurable impact. For example, we included all dogs with heart disease in our ‘exercise intolerance’ category; in moderate to severe cases, this condition will inevitably impact a dog’s ability to exercise (and likely lead to a veterinary recommendation of exercise restriction). However, in mild cases, this condition may have minimal impact on a dog’s ability to exercise.

The most important limitation of our study is that we cannot determine causality given the observational, cross-sectional nature of the design. Given existing knowledge about the relationships between physical activity and cognitive function, it is plausible that higher rates of physical activity play a causal role in reducing risk of later-life cognitive impairment in dogs. However, the observed association between physical activity and cognitive outcomes could also indicate that as dogs decline cognitively, it causes them to become less active. Finally, there is a third possibility of unmeasured confounding, whereby neither physical activity nor cognitive decline have causal effects on one another. The fact that our sensitivity analyses revealed an association between CSLB scores and physical activity even in clinically ‘normal’ dogs suggests that the first explanation is more likely; however, future research incorporating additional study designs, including interventions and the analysis of longitudinal data, will be critical for causal inferences in this domain.

In conclusion, our findings indicate that signs of cognitive decline in dogs, and the likelihood of developing CCD, increase with age. Furthermore, the associations presented here are consistent with the hypothesis that physical activity may partially mitigate these risks, although they are also consistent with the hypothesis that cognitively impaired dogs exercise less, or that unidentified confounding variables influence changes in both physical activity and cognitive function. We also identified several categories of medical conditions that were associated with cognitive dysfunction: sensory deficits showed the strongest associations, and there was also some evidence to suggest associations with endocrine disorders, neurological conditions, orthopedic impairments, periodontal disease, cancer, and kidney disorders. Across a subset of our outcome measures, training history and neuroprotective supplements were associated with reduced cognitive impairment. However, in support of our key hypothesis, physical activity was the only lifestyle factor that was robustly associated with reduced risk of cognitive dysfunction across all three of our outcome measures. These findings establish the value of companion dogs as a model for relationships between physical activity and cognitive aging, and lay a foundation for future longitudinal studies, including randomized controlled trials, with this valuable population.

## Author Contributions

All authors contributed to writing – review & editing. E.B.: conceptualization, methodology, formal analysis, data curation, writing – original draft, and supervision. D.R.: conceptualization and methodology. K.F.: data curation. D.P.: conceptualization, project administration, and funding acquisition. G.A.: conceptualization and methodology. E.M.: conceptualization, methodology, formal analysis, writing – original draft, visualization, and supervision. DAP consortium: resources. G.A. and E.M. both contributed as senior authors.

## Acknowledgments

The Dog Aging Project thanks study participants, their dogs, and community veterinarians for their important contributions.

## Data availability statement

These data are housed on the Terra platform at the Broad Institute of MIT and Harvard.

## Code availability statement

This study did not use custom code or mathematical algorithms.

## Supplementary Information captions

**Supplementary Information 1**. Supplementary tables and appendices.

**Supplementary Information 2**. Summary of HLES items that contributed to each of the following covariates in our full model, along with the total number of unique affected dogs from our sample: sensory impairment, orthopedic, exercise intolerance, neurological, periodontal, liver, endocrine, kidney, and cancer.

**Supplementary Information 3**. A list of all 288 specific health conditions from HLES; Dog Aging Project owners were asked to report, for each condition, whether their dog had been diagnosed. Each of the broad general categories also had an ‘other’ option where owners could write in an answer.

## Dog Aging Project Consortium Authors

Joshua M. Akey^1^, Brooke Benton^2^, Elhanan Borenstein^3,4,5^, Marta G. Castelhano^6^, Amanda E. Coleman^7^, Kate E. Creevy^8^, Kyle Crowder^9,10^, Matthew D. Dunbar^10^, Virginia R. Fajt^11^, Annette L. Fitzpatrick^12,13,14^, Unity Jeffrey^15^, Erica C. Jonlin^2,16^, Matt Kaeberlein^2^, Elinor K. Karlsson^17,18^, Kathleen F. Kerr^19^, Jonathan M. Levine^8^, Jing Ma^20^, Robyn McClelland^19^, Audrey Ruple^21^, Stephen M. Schwartz^13,22^, Sandi Shrager^23^, Noah Snyder-Mackler^24,25,26^, M. Katherine Tolbert^8^, Silvan R. Urfer^2^, Benjamin S. Wilfond^27.28^

^1^Lewis-Sigler Institute for Integrative Genomics, Princeton University, Princeton, NJ, USA

^2^Department of Laboratory Medicine and Pathology, University of Washington School of Medicine, Seattle, WA, USA

^3^Department of Clinical Microbiology and Immunology, Sackler Faculty of Medicine, Tel Aviv University, Tel Aviv, Israel

^4^Blavatnik School of Computer Science, Tel Aviv University, Tel Aviv, Israel

^5^Santa Fe Institute, Santa Fe, NM, USA

^6^Cornell Veterinary Biobank, College of Veterinary Medicine, Cornell University, Ithaca, NY, USA

^7^Department of Small Animal Medicine and Surgery, College of Veterinary Medicine, University of Georgia, Athens, GA, USA

^8^Department of Small Animal Clinical Sciences, Texas A&M University College of Veterinary Medicine & Biomedical Sciences, College Station, TX, USA

^9^Department of Sociology, University of Washington, Seattle, WA, USA

^10^Center for Studies in Demography and Ecology, University of Washington, Seattle, WA, USA

^11^Department of Veterinary Physiology and Pharmacology, Texas A&M University College of Veterinary Medicine & Biomedical Sciences, College Station, TX, USA

^12^Department of Family Medicine, University of Washington, Seattle, WA, USA

^13^Department of Epidemiology, University of Washington, Seattle, WA, USA

^14^Department of Global Health, University of Washington, Seattle, WA, USA

^15^Department of Veterinary Pathobiology, Texas A&M University College of Veterinary Medicine & Biomedical Sciences, College Station, TX, USA

^16^Institute for Stem Cell and Regenerative Medicine, University of Washington, Seattle, WA, USA

^17^Bioinformatics and Integrative Biology, University of Massachusetts Chan Medical School, Worcester, MA, USA

^18^Broad Institute of MIT and Harvard, Cambridge, MA, USA

^19^Department of Biostatistics, University of Washington, Seattle, WA, USA

^20^Division of Public Health Sciences, Fred Hutchinson Cancer Research Center, Seattle, WA, USA

^21^Department of Population Health Sciences, Virginia-Maryland College of Veterinary Medicine, Virginia Tech, Blacksburg, VA, USA

^22^Epidemiology Program, Fred Hutchinson Cancer Research Center, Seattle, WA, USA

^23^Department of Biostatistics, Collaborative Health Studies Coordinating Center, University of Washington, Seattle, WA, USA

^24^School of Life Sciences, Arizona State University, Tempe, AZ, USA

^25^Center for Evolution and Medicine, Arizona State University, Tempe, AZ, USA

^26^School for Human Evolution and Social Change, Arizona State University, Tempe, AZ, USA

^27^Treuman Katz Center for Pediatric Bioethics, Seattle Children’s Research Institute, Seattle, WA, USA

^28^Department of Pediatrics, Divison of Bioethics and Palliative Care, University of Washington School of Medicine, Seattle, WA, USA

