## Supplementary Information 1 for "Associations between physical activity and cognitive dysfunction in older companion dogs: Results from the Dog Aging Project"

**Table S1. Principal components analysis of measures from the Canine**

**Social and Learned Behavior Survey (CSLB).**

| *CSLB item* | *PC1_change* | *PC2_severity* |
| --- | --- | --- |
| CSLB_active_6mo | 0.15 | 0.23 |
| CSLB_defecate_6mo | 0.77 | -0.08 |
| CSLB_food_6mo | 0.87 | 0.04 |
| CSLB_pace_6mo | 0.80 | 0.10 |
| CSLB_recognize_6mo | 0.90 | -0.11 |
| CSLB_stare_6mo | 0.85 | 0.07 |
| CSLB_avoid | -0.03 | 0.38 |
| CSLB_find_food | 0.06 | 0.59 |
| CSLB_pace | -0.05 | 0.68 |
| CSLB_recognize | 0.00 | 0.46 |
| CSLB_stare | 0.01 | 0.69 |
| CSLB_stuck | 0.01 | 0.67 |
| CSLB_walk_walls | 0.02 | 0.64 |
| **Eigenvalue** | **3.55** | **2.58** |
| **Proportion of variance explained** | **0.27** | **0.20** |

*Note: See Appendix A for definitions of each CSLB item.*

**Table S2. Fully adjusted model predicting symptom severity with all dogs, as reported in main results (left column), and when all dogs above the CCD threshold are removed from the analysis (right column). The 95% confidence intervals are based on robust standard errors.**

| Symptom Severity - Sensitivity Analysis | | | | |
| --- | --- | --- | --- | --- |
| **Parameter** | **All Dogs (*n* = 11,574)** | | **Exclude > CCD threshold (*n* = 11,287)** | |
|  | **Beta (95% CI)^1^** | **p-value** | **Beta (95% CI)^1^** | **p-value** |
| physical activity | -0.095 (-0.113 to -0.077) | <0.001 | -0.078 (-0.095 to -0.061) | <0.001 |
| dog weight (lbs) | 0.000 (-0.001 to 0.001) | 0.941 | 0.000 (0.000 to 0.001) | 0.335 |
| sensory impairment | 0.405 (0.349 to 0.461) | <0.001 | 0.320 (0.266 to 0.374) | <0.001 |
| orthopedic | 0.084 (0.041 to 0.127) | <0.001 | 0.079 (0.037 to 0.121) | <0.001 |
| exercise intolerance | 0.043 (-0.023 to 0.108) | 0.203 | 0.047 (-0.017 to 0.111) | 0.153 |
| neurological | 0.073 (0.005 to 0.140) | 0.035 | 0.056 (-0.010 to 0.122) | 0.095 |
| periodontal | 0.060 (0.021 to 0.099) | 0.003 | 0.073 (0.034 to 0.111) | <0.001 |
| liver | 0.030 (-0.193 to 0.253) | 0.790 | -0.001 (-0.216 to 0.213) | 0.990 |
| joint supplement | -0.031 (-0.073 to 0.010) | 0.139 | -0.029 (-0.070 to 0.012) | 0.165 |
| neuroprotective supplement | -0.082 (-0.123 to -0.042) | <0.001 | -0.081 (-0.120 to -0.041) | <0.001 |
| training history | -0.031 (-0.047 to -0.014) | <0.001 | -0.029 (-0.045 to -0.012) | <0.001 |
| endocrine | 0.085 (0.009 to 0.161) | 0.029 | 0.059 (-0.015 to 0.132) | 0.117 |
| kidney | 0.112 (-0.025 to 0.248) | 0.109 | 0.058 (-0.076 to 0.193) | 0.396 |
| cancer | 0.057 (0.001 to 0.113) | 0.047 | 0.056 (0.002 to 0.111) | 0.044 |
| sex |  |  |  |  |
| female intact | — |  | — |  |
| female spayed | 0.195 (0.018 to 0.373) | 0.031 | 0.240 (0.071 to 0.408) | 0.005 |
| male intact | 0.170 (-0.030 to 0.370) | 0.096 | 0.189 (-0.002 to 0.379) | 0.052 |
| male neutered | 0.243 (0.066 to 0.420) | 0.007 | 0.282 (0.113 to 0.450) | 0.001 |
| owner age |  |  |  |  |
| 18-24 | — |  | — |  |
| 25-34 | -0.334 (-0.542 to -0.125) | 0.002 | -0.209 (-0.408 to -0.010) | 0.040 |
| 35-44 | -0.554 (-0.759 to -0.348) | <0.001 | -0.429 (-0.625 to -0.234) | <0.001 |
| 45-54 | -0.707 (-0.911 to -0.503) | <0.001 | -0.587 (-0.781 to -0.393) | <0.001 |
| 55-64 | -0.856 (-1.06 to -0.654) | <0.001 | -0.746 (-0.938 to -0.553) | <0.001 |
| 65-74 | -0.97 (-1.17 to -0.767) | <0.001 | -0.837 (-1.03 to -0.645) | <0.001 |
| 75 and older | -1.05 (-1.25 to -0.839) | <0.001 | -0.915 (-1.11 to -0.716) | <0.001 |
| AIC | 30,003 |  | 28,559 |  |
| ^1^ CI = Confidence Interval | | | | |

**Table S3. Fully adjusted model predicting cognitive decline in previous six months with all dogs, as reported in main results (left column), and when all dogs above the CCD threshold are removed from the analysis (right column). The 95% confidence intervals are based on robust standard errors.**

| Symptom Change - Sensitivity Analysis | | | | |
| --- | --- | --- | --- | --- |
| **Parameter** | **All Dogs (*n* = 11,574)** | | **Exclude > CCD threshold (*n* = 11,287)** | |
|  | **Beta (95% CI)^1^** | **p-value** | **Beta (95% CI)^1^** | **p-value** |
| physical activity | -0.069 (-0.087 to -0.051) | <0.001 | -0.056 (-0.074 to -0.038) | <0.001 |
| dog weight (lbs) | 0.001 (0.000 to 0.002) | 0.004 | 0.001 (0.001 to 0.002) | <0.001 |
| sensory impairment | 0.230 (0.166 to 0.294) | <0.001 | 0.167 (0.104 to 0.229) | <0.001 |
| orthopedic | 0.153 (0.106 to 0.201) | <0.001 | 0.138 (0.092 to 0.185) | <0.001 |
| exercise intolerance | 0.054 (-0.021 to 0.130) | 0.161 | 0.059 (-0.015 to 0.133) | 0.121 |
| neurological | 0.087 (0.011 to 0.163) | 0.025 | 0.076 (0.000 to 0.151) | 0.049 |
| periodontal | 0.063 (0.020 to 0.106) | 0.004 | 0.082 (0.040 to 0.124) | <0.001 |
| liver | -0.023 (-0.298 to 0.253) | 0.872 | -0.039 (-0.320 to 0.242) | 0.783 |
| joint supplement | 0.016 (-0.028 to 0.060) | 0.477 | 0.022 (-0.021 to 0.066) | 0.313 |
| neuroprotective supplement | -0.003 (-0.046 to 0.040) | 0.888 | -0.003 (-0.045 to 0.040) | 0.906 |
| training history | -0.021 (-0.039 to -0.004) | 0.016 | -0.018 (-0.035 to -0.001) | 0.043 |
| endocrine | 0.033 (-0.056 to 0.122) | 0.464 | 0.007 (-0.081 to 0.094) | 0.882 |
| kidney | 0.089 (-0.091 to 0.268) | 0.333 | 0.022 (-0.159 to 0.203) | 0.813 |
| cancer | 0.106 (0.043 to 0.170) | 0.001 | 0.100 (0.038 to 0.161) | 0.002 |
| sex |  |  |  |  |
| female intact | — |  | — |  |
| female spayed | 0.072 (-0.127 to 0.272) | 0.479 | 0.138 (-0.050 to 0.326) | 0.149 |
| male intact | 0.008 (-0.215 to 0.232) | 0.942 | 0.036 (-0.175 to 0.247) | 0.738 |
| male neutered | 0.082 (-0.118 to 0.282) | 0.421 | 0.136 (-0.053 to 0.324) | 0.158 |
| owner age |  |  |  |  |
| 18-24 | — |  | — |  |
| 25-34 | -0.033 (-0.353 to 0.286) | 0.839 | 0.107 (-0.214 to 0.428) | 0.513 |
| 35-44 | -0.060 (-0.378 to 0.258) | 0.713 | 0.084 (-0.236 to 0.403) | 0.607 |
| 45-54 | -0.033 (-0.349 to 0.284) | 0.841 | 0.106 (-0.213 to 0.424) | 0.516 |
| 55-64 | -0.010 (-0.326 to 0.305) | 0.949 | 0.125 (-0.192 to 0.442) | 0.438 |
| 65-74 | -0.077 (-0.393 to 0.238) | 0.632 | 0.076 (-0.240 to 0.393) | 0.636 |
| 75 and older | -0.095 (-0.415 to 0.226) | 0.562 | 0.054 (-0.268 to 0.375) | 0.743 |
| AIC | 31,356 |  | 29,838 |  |
| ^1^ CI = Confidence Interval | | | | |

**Table S4. Fully adjusted models predicting symptom severity for a) all dogs, as reported in main results (left column), b) mixed breed dogs only (middle column), and c) purebred dogs (with at least 10 subjects) only (right column). The Purebreds only model also adjusts for breed. The 95% confidence intervals are based on robust standard errors.**

| Symptom Severity | | | | | | |
| --- | --- | --- | --- | --- | --- | --- |
| **Parameter** | **Full Sample (*n* = 11,574)** | | **Mixed Breed only (*n* = 6,027)** | | **Purebreds only (*n* = 5,167)** | |
|  | **Beta (95% CI)^1^** | **p-value** | **Beta (95% CI)^1^** | **p-value** | **Beta (95% CI)^1^** | **p-value** |
| physical activity | -0.095 (-0.113 to -0.077) | <0.001 | -0.084 (-0.109 to -0.058) | <0.001 | -0.104 (-0.131 to -0.076) | <0.001 |
| dog weight (lbs) | 0.000 (-0.001 to 0.001) | 0.941 | 0.000 (-0.001 to 0.001) | 0.389 | 0.001 (-0.001 to 0.003) | 0.544 |
| sensory impairment | 0.405 (0.349 to 0.461) | <0.001 | 0.368 (0.287 to 0.450) | <0.001 | 0.413 (0.332 to 0.494) | <0.001 |
| orthopedic | 0.084 (0.041 to 0.127) | <0.001 | 0.063 (0.000 to 0.125) | 0.049 | 0.111 (0.048 to 0.174) | <0.001 |
| exercise intolerance | 0.043 (-0.023 to 0.108) | 0.203 | -0.003 (-0.093 to 0.086) | 0.940 | 0.100 (-0.002 to 0.201) | 0.054 |
| neurological | 0.073 (0.005 to 0.140) | 0.035 | 0.046 (-0.054 to 0.147) | 0.366 | 0.100 (0.003 to 0.197) | 0.043 |
| periodontal | 0.060 (0.021 to 0.099) | 0.003 | 0.086 (0.032 to 0.140) | 0.002 | 0.033 (-0.026 to 0.093) | 0.270 |
| liver | 0.030 (-0.193 to 0.253) | 0.790 | -0.119 (-0.425 to 0.186) | 0.443 | 0.224 (-0.103 to 0.550) | 0.179 |
| joint supplement | -0.031 (-0.073 to 0.010) | 0.139 | -0.046 (-0.105 to 0.013) | 0.128 | -0.012 (-0.073 to 0.049) | 0.689 |
| neuroprotective supplement | -0.082 (-0.123 to -0.042) | <0.001 | -0.070 (-0.128 to -0.012) | 0.018 | -0.090 (-0.149 to -0.030) | 0.003 |
| training history | -0.031 (-0.047 to -0.014) | <0.001 | -0.034 (-0.061 to -0.007) | 0.014 | -0.025 (-0.048 to -0.002) | 0.031 |
| endocrine | 0.085 (0.009 to 0.161) | 0.029 | 0.091 (-0.019 to 0.201) | 0.104 | 0.022 (-0.089 to 0.132) | 0.700 |
| kidney | 0.112 (-0.025 to 0.248) | 0.109 | 0.148 (-0.039 to 0.334) | 0.120 | 0.129 (-0.079 to 0.338) | 0.224 |
| cancer | 0.057 (0.001 to 0.113) | 0.047 | 0.053 (-0.024 to 0.130) | 0.177 | 0.074 (-0.011 to 0.159) | 0.089 |
| sex |  |  |  |  |  |  |
| female intact | — |  | — |  | — |  |
| female spayed | 0.195 (0.018 to 0.373) | 0.031 | 0.327 (-0.229 to 0.884) | 0.249 | 0.114 (-0.121 to 0.349) | 0.341 |
| male intact | 0.170 (-0.030 to 0.370) | 0.096 | 0.346 (-0.270 to 0.96) | 0.271 | 0.088 (-0.172 to 0.348) | 0.505 |
| male neutered | 0.243 (0.066 to 0.420) | 0.007 | 0.328 (-0.230 to 0.885) | 0.249 | 0.192 (-0.045 to 0.429) | 0.112 |
| owner age |  |  |  |  |  |  |
| 18-24 | — |  | — |  | — |  |
| 25-34 | -0.334 (-0.542 to -0.125) | 0.002 | -0.289 (-0.528 to -0.050) | 0.018 | -0.364 (-0.807 to 0.080) | 0.108 |
| 35-44 | -0.554 (-0.759 to -0.348) | <0.001 | -0.558 (-0.794 to -0.322) | <0.001 | -0.497 (-0.935 to -0.059) | 0.026 |
| 45-54 | -0.707 (-0.911 to -0.503) | <0.001 | -0.677 (-0.911 to -0.442) | <0.001 | -0.665 (-1.10 to -0.229) | 0.003 |
| 55-64 | -0.856 (-1.06 to -0.654) | <0.001 | -0.847 (-1.08 to -0.615) | <0.001 | -0.781 (-1.22 to -0.346) | <0.001 |
| 65-74 | -0.97 (-1.17 to -0.767) | <0.001 | -0.97 (-1.20 to -0.742) | <0.001 | -0.894 (-1.33 to -0.459) | <0.001 |
| 75 and older | -1.05 (-1.25 to -0.839) | <0.001 | -1.03 (-1.27 to -0.783) | <0.001 | -1.03 (-1.47 to -0.586) | <0.001 |
| ^1^ CI = Confidence Interval | | | | | | |

**Table S5. Fully adjusted models predicting cognitive decline in previous six months for a) all dogs, as reported in main results (left column), b) mixed breed dogs only (middle column), and c) purebred dogs (with at least 10 subjects) only (right column). The Purebreds only model also adjusts for breed. The 95% confidence intervals are based on robust standard errors.**

| Symptom Change: Previous 6 Months | | | | | | |
| --- | --- | --- | --- | --- | --- | --- |
| **Parameter** | **Full Sample (*n* = 11,574)** | | **Mixed Breed only (*n* = 6,027)** | | **Purebreds only (*n* = 5,167)** | |
|  | **Beta (95% CI)^1^** | **p-value** | **Beta (95% CI)^1^** | **p-value** | **Beta (95% CI)^1^** | **p-value** |
| physical activity | -0.069 (-0.087 to -0.051) | <0.001 | -0.058 (-0.084 to -0.032) | <0.001 | -0.095 (-0.123 to -0.066) | <0.001 |
| dog weight (lbs) | 0.001 (0.000 to 0.002) | 0.004 | 0.000 (-0.001 to 0.001) | 0.928 | -0.001 (-0.004 to 0.001) | 0.313 |
| sensory impairment | 0.230 (0.166 to 0.294) | <0.001 | 0.228 (0.136 to 0.320) | <0.001 | 0.202 (0.108 to 0.296) | <0.001 |
| orthopedic | 0.153 (0.106 to 0.201) | <0.001 | 0.183 (0.115 to 0.250) | <0.001 | 0.138 (0.067 to 0.210) | <0.001 |
| exercise intolerance | 0.054 (-0.021 to 0.130) | 0.161 | -0.009 (-0.105 to 0.088) | 0.858 | 0.126 (-0.001 to 0.253) | 0.051 |
| neurological | 0.087 (0.011 to 0.163) | 0.025 | 0.082 (-0.025 to 0.189) | 0.132 | 0.084 (-0.030 to 0.199) | 0.148 |
| periodontal | 0.063 (0.020 to 0.106) | 0.004 | 0.072 (0.013 to 0.130) | 0.017 | 0.087 (0.020 to 0.153) | 0.010 |
| liver | -0.023 (-0.298 to 0.253) | 0.872 | -0.373 (-0.823 to 0.078) | 0.105 | 0.320 (-0.003 to 0.643) | 0.052 |
| joint supplement | 0.016 (-0.028 to 0.060) | 0.477 | 0.057 (-0.005 to 0.118) | 0.072 | -0.030 (-0.098 to 0.038) | 0.386 |
| neuroprotective supplement | -0.003 (-0.046 to 0.040) | 0.888 | -0.033 (-0.093 to 0.027) | 0.284 | 0.023 (-0.042 to 0.089) | 0.489 |
| training history | -0.021 (-0.039 to -0.004) | 0.016 | -0.037 (-0.065 to -0.010) | 0.007 | -0.020 (-0.045 to 0.005) | 0.117 |
| endocrine | 0.033 (-0.056 to 0.122) | 0.464 | 0.080 (-0.048 to 0.209) | 0.222 | -0.021 (-0.147 to 0.106) | 0.751 |
| kidney | 0.089 (-0.091 to 0.268) | 0.333 | 0.171 (-0.046 to 0.389) | 0.123 | 0.043 (-0.269 to 0.356) | 0.786 |
| cancer | 0.106 (0.043 to 0.170) | 0.001 | 0.108 (0.019 to 0.196) | 0.017 | 0.061 (-0.035 to 0.157) | 0.210 |
| sex |  |  |  |  |  |  |
| female intact | — |  | — |  | — |  |
| female spayed | 0.072 (-0.127 to 0.272) | 0.479 | 0.210 (-0.489 to 0.909) | 0.556 | 0.041 (-0.224 to 0.306) | 0.762 |
| male intact | 0.008 (-0.215 to 0.232) | 0.942 | -0.097 (-0.856 to 0.661) | 0.802 | 0.049 (-0.241 to 0.338) | 0.742 |
| male neutered | 0.082 (-0.118 to 0.282) | 0.421 | 0.237 (-0.462 to 0.937) | 0.506 | 0.061 (-0.206 to 0.327) | 0.656 |
| owner age |  |  |  |  |  |  |
| 18-24 | — |  | — |  | — |  |
| 25-34 | -0.033 (-0.353 to 0.286) | 0.839 | -0.101 (-0.454 to 0.252) | 0.576 | 0.225 (-0.493 to 0.942) | 0.539 |
| 35-44 | -0.060 (-0.378 to 0.258) | 0.713 | -0.133 (-0.485 to 0.219) | 0.460 | 0.176 (-0.538 to 0.890) | 0.629 |
| 45-54 | -0.033 (-0.349 to 0.284) | 0.841 | -0.062 (-0.413 to 0.288) | 0.728 | 0.138 (-0.574 to 0.850) | 0.704 |
| 55-64 | -0.010 (-0.326 to 0.305) | 0.949 | -0.064 (-0.413 to 0.285) | 0.718 | 0.190 (-0.520 to 0.901) | 0.600 |
| 65-74 | -0.077 (-0.393 to 0.238) | 0.632 | -0.174 (-0.524 to 0.176) | 0.329 | 0.158 (-0.552 to 0.869) | 0.662 |
| 75 and older | -0.095 (-0.415 to 0.226) | 0.562 | -0.096 (-0.456 to 0.265) | 0.602 | 0.070 (-0.644 to 0.784) | 0.847 |
| ^1^ CI = Confidence Interval | | | | | | |

**Table S6. Fully adjusted models predicting CCD status for a) all dogs, as reported in main results (left column), b) mixed breed dogs only (middle column), and c) purebred dogs (with at least 10 subjects total and one subject with CCD) only (right column). Given the few numbers of dogs with CCD, the Purebreds only model does not adjust for breed.**

| Canine Cognitive Dysfunction (Clinical Cutoff) | | | | | | |
| --- | --- | --- | --- | --- | --- | --- |
| **Parameter** | **Full Sample (*n* = 11,574)** | | **Mixed Breed only (*n* = 6,027)** | | **Purebreds only (*n* = 3,945)** | |
|  | **OR (95% CI)^1^** | **p-value** | **OR (95% CI)^1^** | **p-value** | **OR (95% CI)^1^** | **p-value** |
| physical activity | 0.53 (0.45 to 0.63) | <0.001 | 0.61 (0.47 to 0.77) | <0.001 | 0.47 (0.37 to 0.60) | <0.001 |
| dog weight (lbs) | 0.99 (0.99 to 1.00) | 0.006 | 0.99 (0.98 to 0.99) | 0.002 | 1.00 (0.99 to 1.01) | 0.700 |
| sensory impairment | 3.20 (2.43 to 4.24) | <0.001 | 2.95 (1.95 to 4.51) | <0.001 | 3.16 (2.13 to 4.73) | <0.001 |
| orthopedic | 1.22 (0.92 to 1.62) | 0.162 | 1.26 (0.82 to 1.94) | 0.289 | 1.29 (0.87 to 1.90) | 0.208 |
| exercise intolerance | 0.97 (0.65 to 1.42) | 0.887 | 1.02 (0.55 to 1.80) | 0.943 | 0.79 (0.44 to 1.37) | 0.425 |
| neurological | 1.29 (0.89 to 1.84) | 0.162 | 1.46 (0.83 to 2.49) | 0.176 | 1.43 (0.86 to 2.33) | 0.160 |
| periodontal | 0.78 (0.59 to 1.02) | 0.076 | 0.77 (0.50 to 1.16) | 0.223 | 0.80 (0.54 to 1.18) | 0.273 |
| joint supplement | 1.00 (0.72 to 1.37) | 0.979 | 1.33 (0.82 to 2.14) | 0.241 | 0.79 (0.50 to 1.25) | 0.323 |
| neuroprotective supplement | 0.97 (0.71 to 1.34) | 0.872 | 0.78 (0.49 to 1.26) | 0.317 | 1.34 (0.85 to 2.11) | 0.210 |
| training history | 0.88 (0.74 to 1.03) | 0.133 | 0.96 (0.72 to 1.23) | 0.749 | 0.86 (0.69 to 1.07) | 0.192 |
| endocrine | 1.46 (0.97 to 2.16) | 0.062 | 2.03 (1.13 to 3.49) | 0.013 | 0.91 (0.46 to 1.68) | 0.774 |
| kidney | 1.85 (1.09 to 3.04) | 0.017 | 2.12 (1.02 to 4.18) | 0.036 | 1.66 (0.73 to 3.53) | 0.209 |
| cancer | 1.15 (0.80 to 1.61) | 0.437 | 0.85 (0.47 to 1.46) | 0.576 | 1.24 (0.75 to 2.00) | 0.388 |
| dog sex |  |  |  |  |  |  |
| male | — |  | — |  | — |  |
| female | 0.83 (0.64 to 1.07) | 0.152 | 0.65 (0.44 to 0.96) | 0.031 | 1.05 (0.73 to 1.52) | 0.791 |
| owner age |  |  |  |  |  |  |
| 18-54 | — |  | — |  | — |  |
| 55 and older | 0.78 (0.60 to 1.02) | 0.070 | 0.66 (0.44 to 0.99) | 0.043 | 0.93 (0.64 to 1.38) | 0.719 |
| ^1^ OR = Odds Ratio, CI = Confidence Interval | | | | | | |

**Table S7. Complete list of purebred dogs (*n* = 92) included in sensitivity analyses predicting symptom severity and symptom change, with sample sizes.**

| Breed | Sample size |
| --- | --- |
| Labrador Retriever | 671 |
| Golden Retriever | 505 |
| German Shepherd Dog | 218 |
| Poodle | 188 |
| Dachshund | 165 |
| Australian Shepherd | 156 |
| Border Collie | 139 |
| Chihuahua | 119 |
| Beagle | 110 |
| Shih Tzu | 102 |
| Yorkshire Terrier | 93 |
| Havanese | 84 |
| Miniature Schnauzer | 83 |
| Greyhound | 80 |
| Shetland Sheepdog | 80 |
| Cavalier King Charles Spaniel | 77 |
| Pug | 77 |
| Boston Terrier | 76 |
| West Highland White Terrier | 76 |
| Jack Russell Terrier | 72 |
| Cocker Spaniel | 69 |
| Boxer | 64 |
| Pembroke Welsh Corgi | 64 |
| Siberian Husky | 60 |
| English Springer Spaniel | 57 |
| German Shorthaired Pointer | 57 |
| Australian Cattle Dog | 52 |
| Great Dane | 52 |
| Pomeranian | 52 |
| Doberman Pinscher | 48 |
| American Pitbull Terrier | 47 |
| Cairn Terrier | 47 |
| Poodle (Toy) | 44 |
| Brittany | 43 |
| Soft Coated Wheaten Terrier | 39 |
| Bichon Frise | 38 |
| Weimaraner | 37 |
| Newfoundland | 36 |
| Papillon | 34 |
| American Staffordshire Terrier | 33 |
| Collie | 33 |
| French Bulldog | 32 |
| Maltese | 32 |
| Rat Terrier | 32 |
| Rottweiler | 32 |
| Vizsla | 32 |
| Bulldog | 31 |
| Bernese Mountain Dog | 30 |
| Portuguese Water Dog | 30 |
| Great Pyrenees | 28 |
| Miniature Pinscher | 28 |
| Border Terrier | 27 |
| English Setter | 27 |
| Basset Hound | 26 |
| Cardigan Welsh Corgi | 25 |
| Chesapeake Bay Retriever | 24 |
| Rhodesian Ridgeback | 24 |
| Dalmatian | 21 |
| Miniature American Shepherd | 21 |
| Shiba Inu | 21 |
| Airedale Terrier | 20 |
| Belgian Malinois | 20 |
| Coton De Tulear | 20 |
| Italian Greyhound | 20 |
| Scottish Terrier | 20 |
| Whippet | 20 |
| Samoyed | 19 |
| Alaskan Malamute | 18 |
| American Eskimo Dog | 17 |
| Keeshond | 17 |
| Irish Setter | 16 |
| Lhasa Apso | 15 |
| Belgian Tervuren | 14 |
| Bull Terrier | 14 |
| Catahoula Leopard Dog | 14 |
| English Cocker Spaniel | 14 |
| Mastiff | 14 |
| Old English Sheepdog | 14 |
| Parson Russell Terrier | 14 |
| Pekingese | 14 |
| Flat-Coated Retriever | 13 |
| Welsh Terrier | 13 |
| Carolina Dog | 12 |
| Irish Wolfhound | 12 |
| Wirehaired Pointing Griffon | 12 |
| Treeing Walker Coonhound | 11 |
| Basenji | 10 |
| Bouvier des Flandres | 10 |
| Gordon Setter | 10 |
| Norwich Terrier | 10 |
| Nova Scotia Duck Tolling Retriever | 10 |
| Silky Terrier | 10 |

**Table S8. Complete list of purebred dogs (*n* = 53) included in sensitivity analyses predicting CCD status, with sample sizes.**

| **Breed** | **Sample size** |
| --- | --- |
| Labrador Retriever | 671 |
| Golden Retriever | 505 |
| Poodle | 188 |
| Dachshund | 165 |
| Australian Shepherd | 156 |
| Border Collie | 139 |
| Chihuahua | 119 |
| Beagle | 110 |
| Shih Tzu | 102 |
| Yorkshire Terrier | 93 |
| Havanese | 84 |
| Miniature Schnauzer | 83 |
| Shetland Sheepdog | 80 |
| Cavalier King Charles Spaniel | 77 |
| Pug | 77 |
| Boston Terrier | 76 |
| West Highland White Terrier | 76 |
| Jack Russell Terrier | 72 |
| Cocker Spaniel | 69 |
| Siberian Husky | 60 |
| English Springer Spaniel | 57 |
| German Shorthaired Pointer | 57 |
| Australian Cattle Dog | 52 |
| Pomeranian | 52 |
| Cairn Terrier | 47 |
| Poodle (Toy) | 44 |
| Brittany | 43 |
| Soft Coated Wheaten Terrier | 39 |
| Bichon Frise | 38 |
| Newfoundland | 36 |
| Papillon | 34 |
| French Bulldog | 32 |
| Rat Terrier | 32 |
| Bulldog | 31 |
| Bernese Mountain Dog | 30 |
| Great Pyrenees | 28 |
| Miniature Pinscher | 28 |
| English Setter | 27 |
| Cardigan Welsh Corgi | 25 |
| Shiba Inu | 21 |
| Coton De Tulear | 20 |
| Italian Greyhound | 20 |
| Alaskan Malamute | 18 |
| American Eskimo Dog | 17 |
| Belgian Tervuren | 14 |
| English Cocker Spaniel | 14 |
| Mastiff | 14 |
| Old English Sheepdog | 14 |
| Pekingese | 14 |
| Welsh Terrier | 13 |
| Wirehaired Pointing Griffon | 12 |
| Norwich Terrier | 10 |
| Silky Terrier | 10 |

In our dataset, CSLB scores ranged from 16 to 77, where higher scores indicate higher levels of cognitive dysfunction. However, upon looking closely at the results, we found that scores below 20 occurred when an owner reported both that the dog had no symptoms indicative of cognitive dysfunction, and yet also that the dog had significantly improved over the past 6 months. This scenario seems implausible and is consistent with the owner selecting the left-most response option nearly universally (which corresponds to ‘never’ on the first set of severity questions, but ‘much less’ in response to level of dysfunctional behavior compared to 6 months ago). We therefore removed these 206 dogs from the dataset and re-ran the analyses to ensure that our results did not change.

**Table S9. Fully adjusted models predicting symptom severity for a) the full sample, as reported in the main results (left column) and b) the sample excluding dogs with a CSLB score less than 20 (right column). The 95% confidence intervals are based on robust standard errors.**

| Symptom Severity - Sensitivity Analysis | | | | |
| --- | --- | --- | --- | --- |
| **Parameter** | **Full Sample (*n* = 11,574)** | | **Exclude CSLB < 20 (*n* = 11,368)** | |
|  | **Beta (95% CI)^1^** | **p-value** | **Beta (95% CI)^1^** | **p-value** |
| physical activity | -0.095 (-0.113 to -0.077) | <0.001 | -0.095 (-0.113 to -0.077) | <0.001 |
| dog weight (lbs) | 0.000 (-0.001 to 0.001) | 0.941 | 0.000 (-0.001 to 0.001) | 0.905 |
| sensory impairment | 0.405 (0.349 to 0.461) | <0.001 | 0.404 (0.347 to 0.460) | <0.001 |
| orthopedic | 0.084 (0.041 to 0.127) | <0.001 | 0.084 (0.041 to 0.128) | <0.001 |
| exercise intolerance | 0.043 (-0.023 to 0.108) | 0.203 | 0.048 (-0.018 to 0.114) | 0.153 |
| neurological | 0.073 (0.005 to 0.140) | 0.035 | 0.071 (0.003 to 0.139) | 0.042 |
| periodontal | 0.060 (0.021 to 0.099) | 0.003 | 0.056 (0.017 to 0.096) | 0.005 |
| liver | 0.030 (-0.193 to 0.253) | 0.790 | 0.027 (-0.201 to 0.256) | 0.815 |
| joint supplement | -0.031 (-0.073 to 0.010) | 0.139 | -0.035 (-0.077 to 0.007) | 0.099 |
| neuroprotective supplement | -0.082 (-0.123 to -0.042) | <0.001 | -0.083 (-0.124 to -0.042) | <0.001 |
| training history | -0.031 (-0.047 to -0.014) | <0.001 | -0.029 (-0.045 to -0.012) | <0.001 |
| endocrine | 0.085 (0.009 to 0.161) | 0.029 | 0.095 (0.018 to 0.172) | 0.016 |
| kidney | 0.112 (-0.025 to 0.248) | 0.109 | 0.125 (-0.014 to 0.264) | 0.078 |
| cancer | 0.057 (0.001 to 0.113) | 0.047 | 0.054 (-0.002 to 0.111) | 0.060 |
| sex |  |  |  |  |
| female intact | — |  | — |  |
| female spayed | 0.195 (0.018 to 0.373) | 0.031 | 0.167 (-0.010 to 0.343) | 0.065 |
| male intact | 0.170 (-0.030 to 0.370) | 0.096 | 0.141 (-0.059 to 0.340) | 0.167 |
| male neutered | 0.243 (0.066 to 0.420) | 0.007 | 0.213 (0.036 to 0.390) | 0.018 |
| owner age |  |  |  |  |
| 18-24 | — |  | — |  |
| 25-34 | -0.334 (-0.542 to -0.125) | 0.002 | -0.378 (-0.588 to -0.169) | <0.001 |
| 35-44 | -0.554 (-0.759 to -0.348) | <0.001 | -0.594 (-0.801 to -0.388) | <0.001 |
| 45-54 | -0.707 (-0.911 to -0.503) | <0.001 | -0.751 (-0.96 to -0.546) | <0.001 |
| 55-64 | -0.856 (-1.06 to -0.654) | <0.001 | -0.900 (-1.10 to -0.697) | <0.001 |
| 65-74 | -0.97 (-1.17 to -0.767) | <0.001 | -1.01 (-1.22 to -0.809) | <0.001 |
| 75 and older | -1.05 (-1.25 to -0.839) | <0.001 | -1.09 (-1.30 to -0.882) | <0.001 |
| ^1^ CI = Confidence Interval | | | | |

**Table S10. Fully adjusted models predicting cognitive decline in previous six months for a) the full sample, as reported in the main results (left column) and b) the sample excluding dogs with a CSLB score less than 20 (right column). The 95% confidence intervals are based on robust standard errors.**

| Symptom Severity - Sensitivity Analysis | | | | |
| --- | --- | --- | --- | --- |
| **Parameter** | **Full Sample (*n* = 11,574)** | | **Exclude CSLB < 20 (*n* = 11,368)** | |
|  | **Beta (95% CI)^1^** | **p-value** | **Beta (95% CI)^1^** | **p-value** |
| physical activity | -0.069 (-0.087 to -0.051) | <0.001 | -0.062 (-0.079 to -0.044) | <0.001 |
| dog weight (lbs) | 0.001 (0.000 to 0.002) | 0.004 | 0.001 (0.000 to 0.002) | <0.001 |
| sensory impairment | 0.230 (0.166 to 0.294) | <0.001 | 0.214 (0.151 to 0.277) | <0.001 |
| orthopedic | 0.153 (0.106 to 0.201) | <0.001 | 0.143 (0.097 to 0.189) | <0.001 |
| exercise intolerance | 0.054 (-0.021 to 0.130) | 0.161 | 0.059 (-0.014 to 0.132) | 0.111 |
| neurological | 0.087 (0.011 to 0.163) | 0.025 | 0.086 (0.012 to 0.160) | 0.022 |
| periodontal | 0.063 (0.020 to 0.106) | 0.004 | 0.057 (0.015 to 0.098) | 0.007 |
| liver | -0.023 (-0.298 to 0.253) | 0.872 | -0.003 (-0.276 to 0.270) | 0.984 |
| joint supplement | 0.016 (-0.028 to 0.060) | 0.477 | 0.007 (-0.035 to 0.050) | 0.732 |
| neuroprotective supplement | -0.003 (-0.046 to 0.040) | 0.888 | -0.006 (-0.047 to 0.035) | 0.768 |
| training history | -0.021 (-0.039 to -0.004) | 0.016 | -0.020 (-0.036 to -0.003) | 0.020 |
| endocrine | 0.033 (-0.056 to 0.122) | 0.464 | 0.062 (-0.023 to 0.147) | 0.154 |
| kidney | 0.089 (-0.091 to 0.268) | 0.333 | 0.137 (-0.034 to 0.309) | 0.117 |
| cancer | 0.106 (0.043 to 0.170) | 0.001 | 0.093 (0.031 to 0.155) | 0.003 |
| sex |  |  |  |  |
| female intact | — |  | — |  |
| female spayed | 0.072 (-0.127 to 0.272) | 0.479 | 0.041 (-0.143 to 0.225) | 0.663 |
| male intact | 0.008 (-0.215 to 0.232) | 0.942 | -0.017 (-0.224 to 0.190) | 0.871 |
| male neutered | 0.082 (-0.118 to 0.282) | 0.421 | 0.042 (-0.142 to 0.227) | 0.653 |
| owner age |  |  |  |  |
| 18-24 | — |  | — |  |
| 25-34 | -0.033 (-0.353 to 0.286) | 0.839 | -0.131 (-0.438 to 0.177) | 0.404 |
| 35-44 | -0.060 (-0.378 to 0.258) | 0.713 | -0.141 (-0.447 to 0.165) | 0.366 |
| 45-54 | -0.033 (-0.349 to 0.284) | 0.841 | -0.118 (-0.422 to 0.187) | 0.448 |
| 55-64 | -0.010 (-0.326 to 0.305) | 0.949 | -0.078 (-0.381 to 0.225) | 0.616 |
| 65-74 | -0.077 (-0.393 to 0.238) | 0.632 | -0.125 (-0.428 to 0.178) | 0.420 |
| 75 and older | -0.095 (-0.415 to 0.226) | 0.562 | -0.156 (-0.464 to 0.152) | 0.320 |
| ^1^ CI = Confidence Interval | | | | |

**Table S11. Fully adjusted models predicting CCD status for a) the full sample, as reported in the main results (left column) and b) the sample excluding dogs with a CSLB score less than 20 (right column).**

| Canine Cognitive Dysfunction (Clinical Cutoff) – Sensitivity Analysis | | | | |
| --- | --- | --- | --- | --- |
| **Parameter** | **Full Sample (*n* = 11,574)** | | **Exclude CSLB < 20 (*n* = 11,368)** | |
|  | **OR (95% CI)^1^** | **p-value** | **OR (95% CI)^1^** | **p-value** |
| physical activity | 0.53 (0.45 to 0.63) | <0.001 | 0.54 (0.45 to 0.63) | <0.001 |
| dog weight (lbs) | 0.99 (0.99 to 1.00) | 0.006 | 0.99 (0.99 to 1.00) | 0.006 |
| sensory impairment | 3.20 (2.43 to 4.24) | <0.001 | 3.17 (2.40 to 4.20) | <0.001 |
| orthopedic | 1.22 (0.92 to 1.62) | 0.162 | 1.22 (0.92 to 1.61) | 0.165 |
| exercise intolerance | 0.97 (0.65 to 1.42) | 0.887 | 0.98 (0.65 to 1.43) | 0.906 |
| neurological | 1.29 (0.89 to 1.84) | 0.162 | 1.29 (0.89 to 1.84) | 0.163 |
| periodontal | 0.78 (0.59 to 1.02) | 0.076 | 0.78 (0.59 to 1.02) | 0.073 |
| joint supplement | 1.00 (0.72 to 1.37) | 0.979 | 0.99 (0.72 to 1.37) | 0.971 |
| neuroprotective supplement | 0.97 (0.71 to 1.34) | 0.872 | 0.97 (0.71 to 1.34) | 0.863 |
| training history | 0.88 (0.74 to 1.03) | 0.133 | 0.88 (0.75 to 1.03) | 0.139 |
| endocrine | 1.46 (0.97 to 2.16) | 0.062 | 1.48 (0.98 to 2.18) | 0.057 |
| kidney | 1.85 (1.09 to 3.04) | 0.017 | 1.88 (1.11 to 3.08) | 0.016 |
| cancer | 1.15 (0.80 to 1.61) | 0.437 | 1.15 (0.81 to 1.62) | 0.425 |
| dog sex |  |  |  |  |
| male | — |  | — |  |
| female | 0.83 (0.64 to 1.07) | 0.152 | 0.83 (0.64 to 1.08) | 0.163 |
| owner age |  |  |  |  |
| 18-54 | — |  | — |  |
| 55 and older | 0.78 (0.60 to 1.02) | 0.070 | 0.79 (0.61 to 1.03) | 0.078 |
| ^1^ OR = Odds Ratio, CI = Confidence Interval | | | | |

**Fig S1.** Bivariate relationship between age and physical activity with 95% confidence intervals (grey shading).

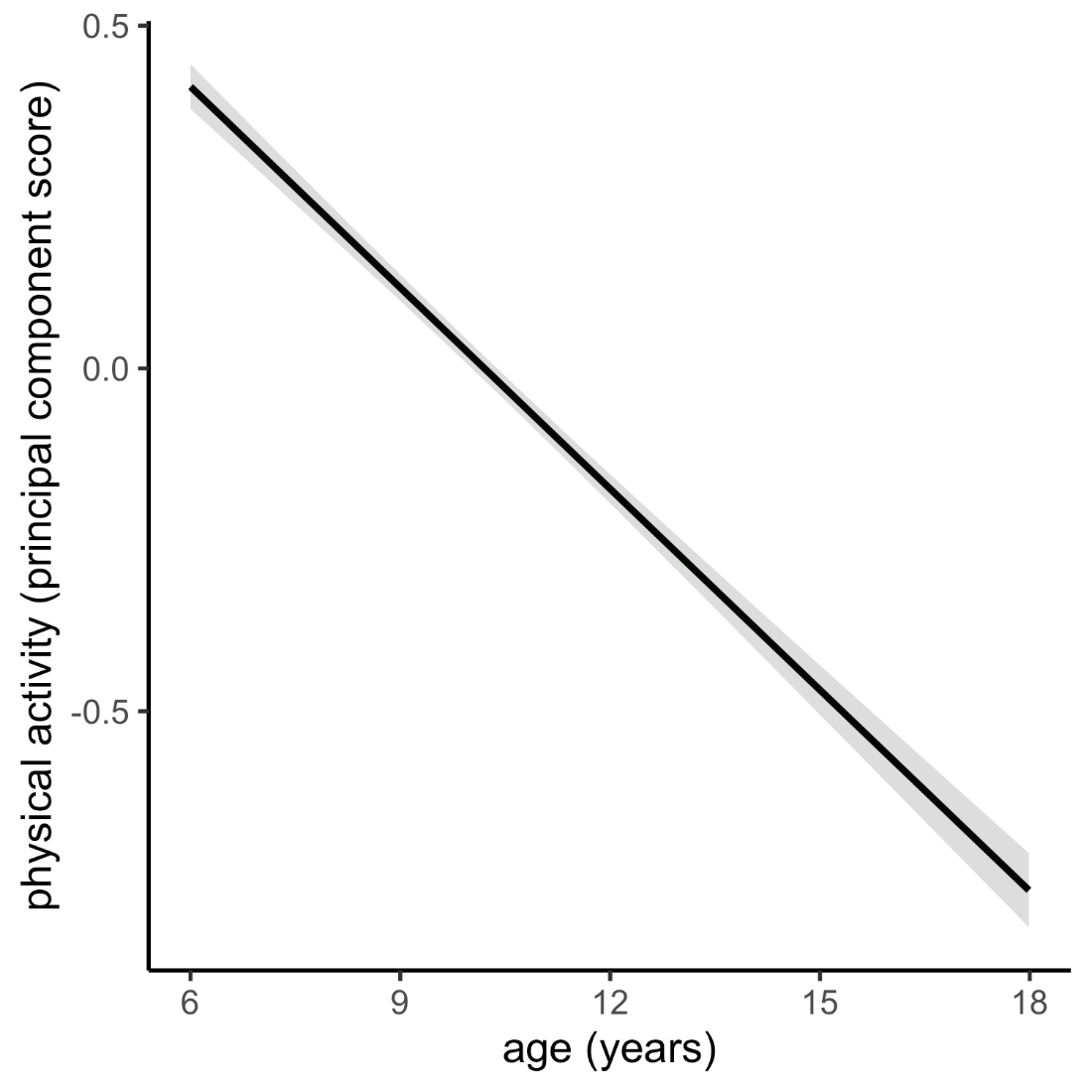

**Fig S2.** Plots show the fitted values and 95% confidence intervals (grey shading) between age and the cognitive measures: severity (left), change (middle), and CCD status (right). Plots in the top row reflect a fitted effect using a linear term for age, whereas the plots in the bottom row use a second order polynomial for age. For the symptom severity and change panels, the points are the mean ± SEM at each age year. For the CCD status panels, the points are the proportion of dogs with CCD.

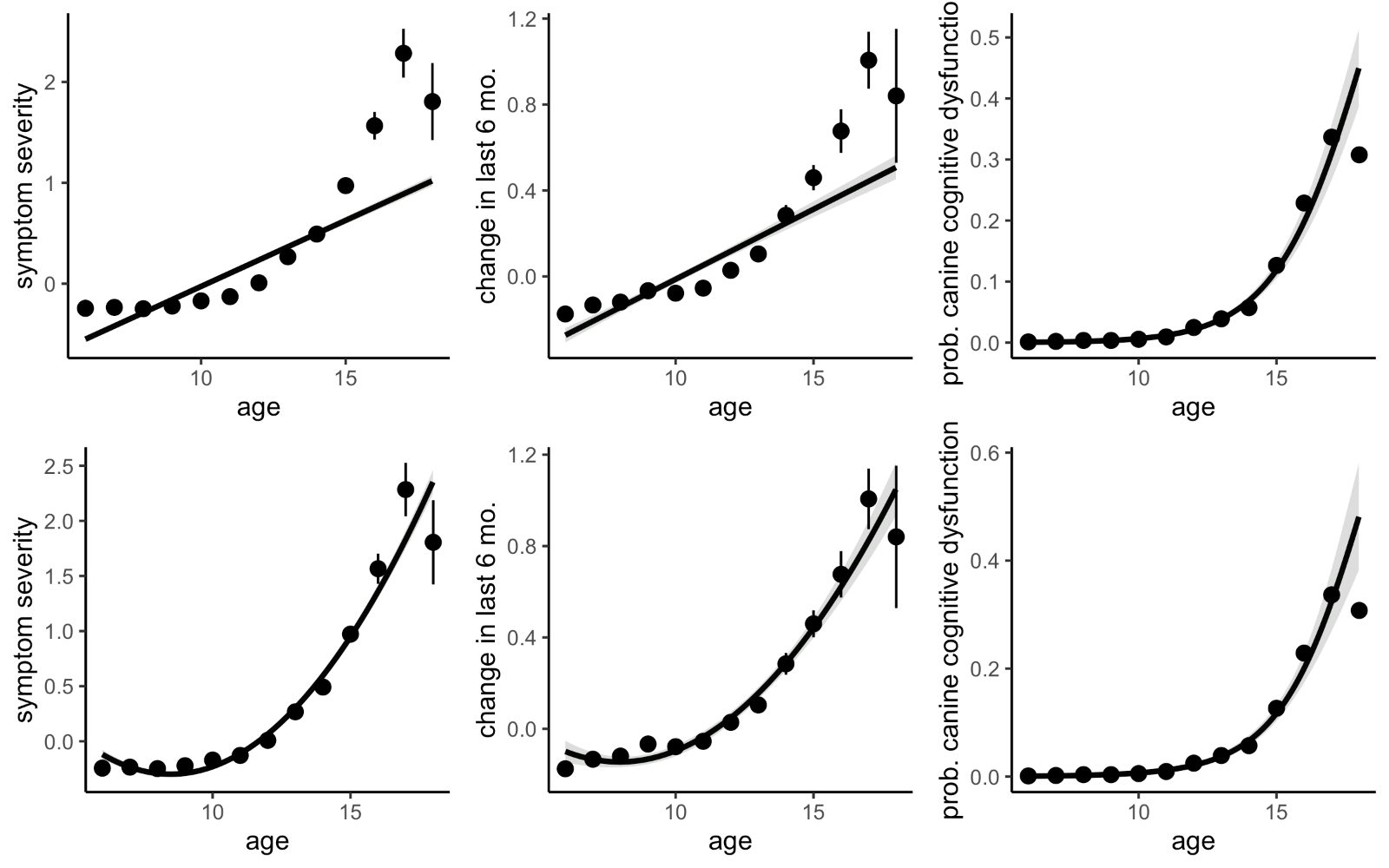

**Appendix A.**

**Dog Aging Project: Canine Social and Learned Behavior survey.**

The goal of this survey is to assess age‐related cognitive and behavioral changes in dogs. The ﬁrst time you ﬁll it out establishes a baseline score for your dog. You will have the opportunity to complete the survey again every year. This repeated design allows us to learn how the dogs in the study change over time.

None of the behaviors described in the survey are bad or wrong. These behaviors are exhibited at various times by many kinds of dogs, and they give our researchers a glimpse into underlying cognitive processes. This speciﬁc assessment tool was developed and validated by Dr. Hannah Salvin (full reference below).

Answers to all questions are required. Please try to answer to the best of your ability based on your dog at their current age.

Salvin HE et al (2011). CCDR: A data‐driven and ecologically relevant assessment tool. *The Veterinary Journal* 188: 331‐336.

Using the descriptions provided, please select the option that best matches your dog.

Never (1)

Once a month (2)

Once a week (3)

Once a day (4)

More than once a day (5)

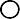

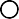

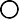

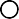

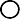
How often does your dog pace up and down, walk in circles and/or wander with no direction or purpose?

**[cslb_pace]**

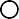

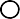

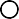

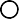

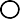
How often does your dog stare blankly at the walls

or ﬂoor?

**[cslb_stare]**

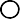

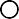

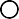

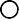

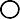
How often does your dog get stuck behind objects

and is unable to get around?

**[cslb_stuck]**

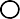

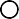

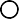

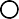

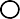
How often does your dog fail to recognize familiar

people or other pets?

**[cslb_recognize]**

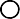

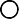

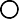

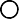

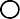
How often does your dog walk into walls or doors?

**[cslb_walk_walls]**

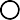

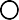

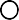

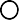

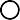
How often does your dog walk away while, or avoid,

being petted?

**[cslb_avoid]**

Using the descriptions provided, please select the option that best matches your dog.

Never (1)

1‐30% of the time (2)

31‐60% of the time (3)

61‐99% of

the time (4)	Always (5)

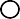

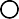

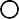

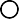

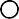
How often does your dog have diﬃculty ﬁnding food dropped on the ﬂoor? **[cslb_ﬁnd_food]**

Using the descriptions listed above each box, place a checkmark in the box that best matches your dog. If your dog has never exhibited the behavior, check "The same."

Much less (1)

Slightly	less (2)

The same	(3)

Slightly more (4)

Much more (5)

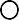

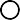

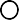

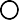

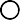
Compared with 6 months ago, how much does your dog now pace up and down, walk in circles and/or wander

with no direction or purpose? **[cslb_pace_6mo]**

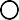

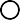

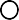

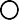

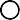
Compared with 6 months ago, how much does your dog

now stare blankly at the walls or ﬂoor? **[cslb_stare_6mo]**

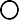

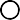

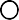

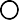

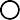
Compared with 6 months ago, how much does your dog urinate or defecate in an area it has previously kept clean?

**[cslb_defecate_6mo]**

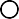

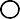

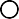

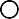

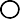
Compared with 6 months ago, how much does your dog have diﬃculty ﬁnding food dropped on the ﬂoor?

**[cslb_food_6mo]**

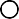

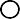

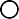

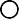

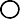
Compared with 6 months ago, how much does your dog fail to recognize familiar people or other pets?

**[cslb_recognize_6mo]**

Using the descriptions listed above each box, place a checkmark in the box that best matches your dog. If your dog has never exhibited the behavior, check "The same."

Much

more (1)

Slightly more (2)

The same (3)

Slightly less (4)

Much

less (5)

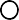

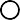

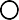

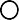

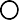
Compared with 6 months ago, how much time does

your dog spend active? **[cslb_active_6mo]**

*Optional:* In the last 6 months, have you observed any other changes in your dog’s thinking or information

processing that you want to share with us? **[cslb_other_changes]**

**Woof! Thank you for completing the Canine Social and Learned Behavior Survey!**

**Calculating CSLB Score**

- The value next to each item indicates how many points are given for a response. For example, a response of “once a month (2)” to the ﬁrst item is scored as 2 points.
- Multiply the response for “Compared with 6 months ago, how much does your dog have diﬃculty ﬁnding food dropped on the ﬂoor?” by 2 (maximum 10 points)
- Multiply the response for “Compared with 6 months ago, how much does your dog fail to recognize familiar people or other pets?” by 3 (maximum 15 points)
- Add all points together (including items that were multiplied) to determine the CSLB score.

**CSLB Score:** _**[cslb_score**_**]**

Threshold >= 50 is diagnostic of canine cognitive dysfunction

**Derived variable [cslb_score] = [cslb_pace] + [cslb_stare] + [cslb_stuck] + [cslb_recognize] + [cslb_walk_walls] + [cslb_avoid] + [cslb_ﬁnd_food] + [cslb_pace_6mo] + [cslb_stare_6mo] + [cslb_defecate_6mo] + (2 * [cslb_food_6mo]) + (3 * [cslb_recognize_6mo]) + [cslb_active_6mo]**

**Appendix B.**

Below we provide the detailed criteria for whether a dog had a history of training (coded as a binary variable), determined by considering HLES responses for a dog’s ‘Primary activity’ and ‘Secondary activity’.

**If only one activity was reported:**

| **Category** | **History of training (1/0)** | **Justification** |
| --- | --- | --- |
| Companion animal or pet | 0 | No explicit training required |
| Service dog (Seeing eye dog, Hearing or signal dog, Wheelchair service dog) | 1 | Requires training |
| Assistance or therapy dog | Varies | See subcategories below |
| Obedience | 1 | Requires training |
| Agility | 1 | Requires training |
| Working (herding, guarding, etc.) | 1 | Requires training |
| Hunting | 1 | Requires training |
| Show | 1 | Requires training |
| Search & Rescue | 1 | Requires training |
| Field Trials | 1 | Requires training |
| Breeding | 0 | No explicit training required |
| Other | Varies | See guidelines below |

**Assistance or therapy dog subcategories:**

| **Category** | **History of training (1/0)** | **Justification** |
| --- | --- | --- |
| Community therapy dog | 1 | Requires training |
| Emotional support dog | 0 | No explicit training required |

**Guidelines for write-in answers from ‘other’ categories (including Other, Other medical service dog, Other health assistance dog):**

| **Category** | **History of training (1/0)** | **Justification** |
| --- | --- | --- |
| Not enough information (e.g., demonstration dog but no indication of for what) | Excluded | Unclear if training is involved |
| Athlete | Excluded | Unclear if training is involved or just recreational |
| Mobility support | 1 | Requires training |
| PTSD service dog | 1 | Requires training |
| Facility dog (e.g., courtroom dog) | 1 | Requires training |
| Psychiatric service dog | 1 | Requires training |
| Medical alert dog (e.g., diabetic alert) | 1 | Requires training |
| Nose work/scent work | 1 | Requires training |
| Police K9 | 1 | Requires training |
| Sled dog mushing | 1 | Requires training |
| Mentions training or obedience work | 1 | Requires training |
| Autism service dog | 1 | Requires training |
| Greyhound/track racing | 1 | Requires training |
| Flyball | 1 | Requires training |
| Dock diving | 1 | Requires training |
| Disc | 1 | Requires training |
| Rally | 1 | Requires training |
| Barn Hunt | 1 | Requires training |
| Frisbee | 1 | Requires training |
| Reading therapy | 0 | Unless specifically designated as a 'therapy dog' or as a participant in a recognized therapy dog program; otherwise: no explicit training required |
| Visits to hospitals/schools/nursing homes | 0 | Unless specifically designated as a 'therapy dog'; otherwise: no explicit training required |
| Blood donor | 0 | No explicit training required |
| Exercise/hiking/joring/walking buddy | 0 | No explicit training required |
| Whimsical, tongue-in-cheek responses (e.g., keeps me smiling) | 0 | No explicit training required |
| Office dog/greeter | 0 | No explicit training required |
| Guard dog/Watch dog | 0 | No explicit training required |
| Unofficial/"Untrained" therapy dog | 0 | No explicit training required |
| Comfort dog / Mental Support | 0 | No explicit training required |
| Distraction from pain | 0 | No explicit training required |
| Lure Coursing | 0 | No explicit training required |

**Guidelines for dogs with both a primary and secondary activity:**

- If at least one activity was determined to involve training, the dog was coded as having a history of training (1)
- If both activities were determined to not need explicit training, the dog was coded as *not* having a history of training (0)
- If one activity was determined to not need explicit training and the other was unable to be determined, we were unable to determine the ‘history of training’ score and these dogs were excluded from the CSLB analysis
